# Cellular and molecular phenotypes of *C9orf72* ALS/FTD patient derived iPSC-microglia mono-cultures

**DOI:** 10.1101/2020.09.03.277459

**Authors:** Ileana Lorenzini, Eric Alsop, Jennifer Levy, Lauren M Gittings, Deepti Lall, Benjamin E Rabichow, Stephen Moore, Ryan Pevey, Lynette Bustos, Camelia Burciu, Divya Bhatia, Mo Singer, Justin Saul, Amanda McQuade, Makis Tzioras, Thomas A Mota, Amber Logemann, Jamie Rose, Sandra Almeida, Fen-Biao Gao, Michael Marks, Christopher J Donnelly, Elizabeth Hutchins, Shu-Ting Hung, Justin Ichida, Robert Bowser, Tara Spires-Jones, Mathew Blurton-Jones, Tania F Gendron, Robert H Baloh, Kendall Van Keuren-Jensen, Rita Sattler

## Abstract

While motor and cortical neurons are affected in *C9orf72* ALS/FTD, it remains still largely unknown if and how non-neuronal cells induce or exacerbate neuronal damage. We generated *C9orf72* ALS/FTD patient-derived induced pluripotent stem cells differentiated into microglia (iPSC-MG) and examined their intrinsic phenotypes. Similar to iPSC motor neurons, *C9orf72* ALS/FTD iPSC-MG mono-cultures form G_4_C_2_ repeat RNA foci, exhibit reduced C9orf72 protein levels and generate dipeptide repeat proteins. Healthy control and *C9orf72* iPSC-MG equivalently express microglial specific genes and display microglial functions including inflammatory cytokine release and phagocytosis of extracellular toxic cargos such as synthetic amyloid beta peptides and healthy human brain synaptoneurosomes. Select *C9orf72* iPSC-MG patient lines show inability to efficiently remove phagocytosed contents, suggesting dysfunction of the endosomal-lysosomal pathways. Finally, RNA sequencing revealed overall transcriptional changes in diseased microglia yet no significant differentially expressed microglial-enriched genes. These minimal differences in cellular, molecular and functional characteristics of microglial mono-cultures suggest that a diseased microenvironment is associated with microglial activation and subsequent regulation of neuronal dysfunction.

## INTRODUCTION

The GGGGCC (G_4_C_2_) hexanucleotide repeat expansion (HRE) in the non-coding region of the *chromosome 9 open reading frame 72* (*C9orf72)* gene is considered the most prevalent genetic abnormality associated with the spectrum disease of amyotrophic lateral sclerosis and frontotemporal dementia (ALS/FTD) to date (DeJesus-Hernandez et al., 2011, Renton et al., 2011b). The *C9orf72* HRE has been hypothesized to contribute to neurodegeneration through three non-mutually exclusive mechanisms. First, *C9orf72* HRE leads to haploinsufficiency and reduced C9orf72 protein expression due to a failure in transcription of the expanded allele; second, non-canonical translation of the repeat RNA leads to synthesis of dipeptide repeat (DPR) proteins; third, toxic RNA gain-of-function is suggested via sequestration of RNA binding proteins to G_4_C_2_ RNA foci. Additionally, *C9orf72* postmortem tissues exhibit TAR-DNA binding protein 43 (TDP-43) pathology, characterized by nuclear depletion and cytoplasmic inclusions of TDP-43 (Josephs et al., 2016, Neumann et al., 2006, Ling et al., 2013, Lagier-Tourenne et al., 2010, Nakashima-Yasuda et al., 2007, Uryu et al., 2008).

While neuronal degeneration is a hallmark of ALS/FTD, it is well known that glia can impact the onset and progression of diseases via non-cell autonomous disease mechanisms (Ilieva et al., 2009, Yamanaka et al., 2008, Haidet-Phillips et al., 2011, Barbeito et al., 2004, Brites and Vaz, 2014, Frakes et al., 2014, Yamanaka and Yamashita, 2007, Kang et al., 2013, Madill et al., 2017, Zhao et al., 2020, Ghasemi et al., 2021). *C9orf72,* known to be differentially expressed in the central nervous system (CNS), has its highest expression observed in peripheral myeloid cells and microglia (Rizzu et al., 2016, O’Rourke et al., 2016, Zhang et al., 2016a), stressing the need to understand the role and contribution of mutant *C9orf72* microglia to disease pathogenesis. *C9orf72* ALS/FTD postmortem brain tissues exhibit extensive microglial pathology in the corticospinal tract and corpus callosum compared to non-*C9orf72* ALS patients and control cases (Brettschneider et al., 2012, Cooper-Knock et al., 2012, Cardenas et al., 2017). Other reports showed enlarged CD68-positive lysosomes in microglia of the motor cortex and spinal cord of *C9orf72*-ALS patients compared to sporadic ALS (sALS), indicating active phagocytic activity in these regions (O’Rourke et al., 2016). With respect to *C9orf72* ALS/FTD cellular phenotypes, it is notable that pathological features examined in postmortem autopsy tissues, such as the formation of nuclear RNA foci and the generation of DPR proteins, are most prominent in neurons compared to neighboring astrocytes and microglia (Rostalski et al., 2019, Mizielinska et al., 2013, Mackenzie et al., 2013, Saberi et al., 2018, DeJesus-Hernandez et al., 2017, Gendron et al., 2013).

While several *C9orf72* ALS/FTD mouse models exhibit gliosis and inflammation (O’Rourke et al., 2016, Liu et al., 2016, Schludi et al., 2017, Zhang et al., 2018), mouse models of *C9orf72*-deficiency show a more defined contribution of microglia to disease phenotypes. *C9orf72* knockout mice display microglia lysosomal accumulation and increased expression of pro-inflammatory cytokines including IL-6 and IL-1β (O’Rourke et al., 2016, Jiang et al., 2016, Burberry et al., 2016). In addition, transcriptomic profiling revealed an upregulation of inflammatory pathways similar to what has been found in *C9orf72* FTD patient tissues (Prudencio et al., 2015, O’Rourke et al., 2016). Likewise, knocking out *C9orf72* with antisense oligonucleotides in mice led to the upregulation of *TREM2* and *C1qa,* which are both upregulated in activated microglia (Lagier-Tourenne et al., 2013a). Furthermore, we recently showed that *C9orf72*-deficient microglia change their transcriptional profile to an enhanced inflammatory type I IFN signature and promote microglia-mediated synapse loss, suggesting a direct contribution of these cells to neurodegeneration (Lall et al., 2021). While these *in vivo* mouse models indicate that loss of *C9orf72* in microglia displays non-cell autonomous regulatory activities, it is still unknown whether the presence of endogenous *C9orf72* HREs in patient-derived human microglia leads to inherent phenotypic changes, which in turn could contribute non-cell autonomously to *C9orf72* disease pathogenesis. Similarly, microglia responses and activation states are known to be unique to the disease environment and therefore, neuron-microglia communication may be required to regulate mutant *C9orf72* microglial activation and exacerbation of disease phenotypes during neurodegeneration in *C9orf72* ALS/FTD. This hypothesis is supported by recent studies using microglia single-cell expression patterns to show that while there is a conserved gene core program, a spectrum of microglia responses and activation states are unique to the disease environment (Galatro et al., 2017, Smith and Dragunow, 2014, Gosselin et al., 2017, Geirsdottir et al., 2019, Friedman et al., 2018). Finally*, in vitro* studies on intrinsic properties of *C9orf72* microglia were recently assessed by overexpressing *C9orf72* HRE in mouse BV-2 microglial cells (Rostalski et al., 2020). While BV-2 microglial cells presented with *C9orf72-*associated phenotypes, such as production of DPR proteins, the presence of *C9orf72* HRE in microglia did not alter their functions or viability.

To better understand the contribution of microglial cells to *C9orf72* ALS/FTD, we used an endogenous human *in vitro* cell culture model by differentiating *C9orf72* ALS/FTD patient derived induced pluripotent stem cell into microglia (iPSC-MG). These iPSC-MG mono-cultures express known microglial genes and proteins. iPSC-MG carrying the endogenous *C9orf72* HRE recapitulate aspects of *C9orf72* ALS/FTD disease pathogenesis such as the formation of HRE-associated RNA foci, expression of poly (GP) DPRs and reduced C9orf72 protein levels due to *C9orf72* haploinsufficiency, which was corrected in isogenic control iPSC-MG and not present in healthy control iPSC-MGs. Interestingly, despite the presence of this *C9orf72* pathobiology, iPSC-MG grown in mono-cultures show minor gene expression changes compared to control iPSC-MG. Similarly, *C9orf72* and control iPSC-MG perform common microglial functions such as the release of cytokines and chemokines upon exposure to extracellular lipopolysaccharide (LPS) stimuli. In addition, *C9orf72* iPSC-MG showed cytochalasin-D dependent phagocytic activity upon human healthy brain synaptoneurosomes exposure and engulfed amyloid beta (Aβ) (1-40)-TAMRA indistinguishable from healthy control iPSC-MG. Select *C9orf72* patient lines did exhibit increased retention of engulfed toxic cargo, suggesting patient-specific alterations of the endosomal and lysosomal pathway. These studies provide the first characterization of the cellular and molecular phenotypes of endogenous *C9orf72* ALS/FTD patient derived iPSC-MG mono-cultures. Our data suggest that a diseased CNS microenvironment contributes to the disease-specific microglial phenotypic changes observed in postmortem patient brain tissues.

## RESULTS

### *C9orf72* ALS/FTD patient and control iPSCs differentiate into brain-like microglia

Following established protocols (McQuade et al., 2018, Abud et al., 2017) we differentiated control and *C9orf72* ALS/FTD patient iPSC lines into microglia (Figure 1A; Table S1-S3). At a mature stage of 40 days *in vitro* (DIV), both control and *C9orf72* ALS/FTD iPSC-MG display a typical ramified microglia morphology (Figure 1B) and express classic microglial marker proteins including the myeloid transcription factor PU.1, purinergic surface receptor P2RY12 and C-X3-C Motif Chemokine Receptor 1 (CX3CR1), as confirmed via fluorescent immunocytochemistry (Figure 1C; Figures S1A). Differentiated microglia uniformly expressed the triggering receptor expressed on myeloid cells 2 (TREM2) and transmembrane protein 119 (TMEM119), further demonstrating a commitment to microglial fate, with no differences detected between *C9orf72* and control iPSC-MG (Figure 1C; Figures S1A and S1B). To validate microglial lineage, we performed Illumina paired-end deep RNA sequencing analysis (RNA seq) on control and *C9orf72* ALS/FTD iPSC-MG (Figures 1D and 1E). At a transcriptional level, control and *C9orf72* iPSC-MG clustered together and presented with a unique transcriptome compared to iPSCs differentiated into forebrain cortical neurons (iPSC-CN) (Figure 1D). Principal component analysis of this RNA seq dataset revealed a highly similar gene expression profile within iPSC-MG and different from iPSC-CNs confirming that these populations are distinct from each other (PC1, 94.87% variance; PC2, 1.66% variance; Figure 1E). Moreover, to verify the absence of other cell types in the differentiated iPSC-MG cultures, we quantified known cell type specific markers revealing high expression of microglial-enriched genes and low expression for transcripts unique to astrocytes, oligodendrocyte precursor cells (OPC), oligodendrocytes, neurons, endothelial cells and pericytes (Zhang et al., 2014). We observed expression of endothelial and pericyte marker genes *FTL1, GPC3* and *FMOD*, however these genes are known to also be expressed in microglia (https://www.proteinatlas.org/) (Figure 1F). These data confirm that the differentiation of microglia from iPSC results in brain-like microglia that express microglial-enriched genes and proteins, distinct from other cell types, yet no inherent differences were observed between control and human *C9orf72* ALS/FTD patient iPSC-MGs.

**Figure 1.**
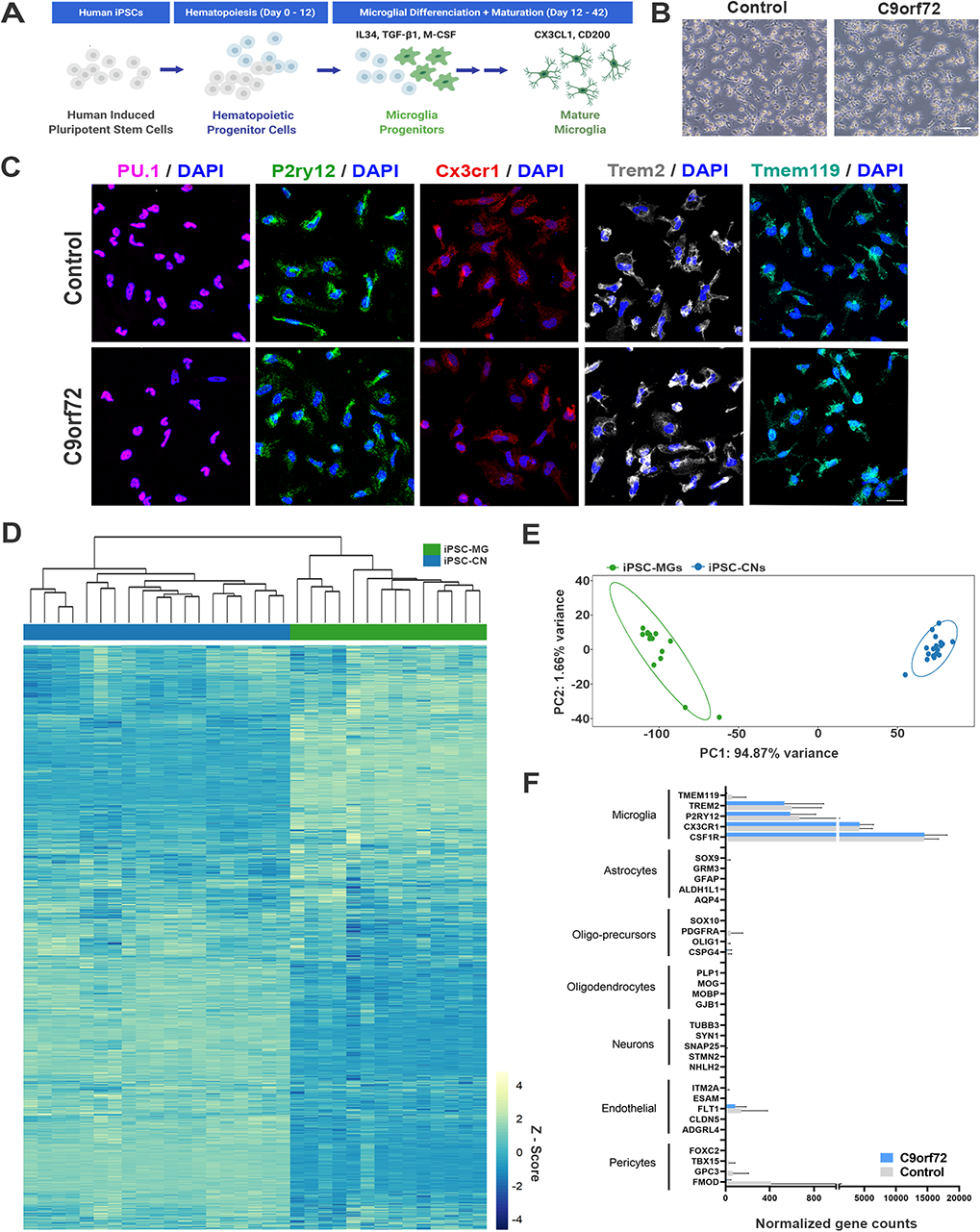
Healthy control and *C9orf72* ALS/FTD patient iPSC lines differentiate into mature microglia. Healthy control and *C9orf72* ALS/FTD iPSC-MG monocultures were generated and examined for classic brain microglia characteristics. (A) Schematic illustration of iPSC-MG differentiation protocol (adapted from (McQuade et al., 2018, Abud et al., 2017)). (B) Phase contrast images of mature iPSC-MG differentiated from healthy control and *C9orf72* ALS/FTD iPSCs (DIV 40). The representative images show typical ramified microglia morphology in both experimental groups. Scale bar, 120μm. (C) Representative immunofluorescence of DIV 40 control (n=4) and *C9orf72* ALS/FTD iPSC-MG (n=7) stained for myeloid transcription factor PU.1 and microglia specific markers such as purinergic surface receptor P2ry12, C-X3-C Motif Chemokine Receptor 1 (Cx3cr1), triggering receptor expressed on myeloid cells 2 (TREM2) and the transmembrane protein 119 (TMEM119). For quantification of marker protein expression/percentage of DAPI-positive cells, see Figure S1A. For quantitative marker gene expression levels, see Figure S1B. Scale bar, 20μm. (D) Heatmap of the complete iPSC-MG (control, n=4 cell lines with 1-2 differentiations each and *C9orf72* ALS/FTD, n=7 cell lines with 1-2 differentiations each) and iPSC-CN (n=12 cell lines with 1-3 differentiations each) transcriptome demonstrating distinct gene expression profiles between the two cell populations. All iPSC-MG and iPSC-CN samples were normalized together by DESeq2 and Z-score scaled. (E) Principal component analysis of the RNA-seq expression data revealed a highly similar gene expression profile within both iPSC-MG and iPSC-CNs (green cluster) and confirmed these populations as distinct from each other (blue cluster) (PC1, 94.87% variance; PC2, 1.66% variance). (F) Normalized counts for genes associated with microglia, astrocytes, oligo-precursor cells (OPC), oligodendrocytes, neurons, endothelial cells and pericytes within the iPSC-MG population (Control n=4 lines; *C9orf72* ALS/FTD n=7 lines). Gene list from (Zhang et al., 2014). *FTL1, GPC3* and *FMOD*, endothelial and pericyte marker genes known to be expressed in microglia (https://www.proteinatlas.org/). Bar graphs are presented as mean ± SD. Heatmaps were generated using z-scores calculated from DESeq2 normalized gene counts. Principal component analysis was done using DESeq2 normalized gene counts with the PCAExplorer package (v. 2.14.1) in R. Bar graphs show normalized gene counts per iPSC-MG group generated from the RNA sequencing presented as Mean ± SD.

### The transcriptional profile of microglial-enriched genes in *C9orf72* ALS/FTD iPSC-MG resembles that of control iPSC-MG

To test if *C9orf72* microglia exhibit intrinsic disease-mediated gene expression alterations, we analyzed our RNA sequencing data for overall differential expression between mature control and *C9orf72* ALS/FTD iPSC-MG. This analysis revealed several differently expressed genes (DEG) with significantly altered expression in *C9orf72* ALS/FTD iPSC-MG (20 genes; log_2_ fold change (FC) ± 1, p value <0.05; Figure 2A; Figure S2A). The topmost dysregulated genes included *HOGA1*, *SEC14L5*, *FPR3*, *HSPA2*, *KCNK17*, *L1TD1*, *CLEC12A* and *VAV3,* some of which have been described to be associated with neuroinflammation and neurodegeneration (Busch et al., 2022, Leak, 2014, Petyuk et al., 2018). We next determined if 881 RNA transcripts previously reported to be enriched in cortical microglia were differentially expressed in our dataset but found no significant dysregulation of these select genes (Figure 2B) (Gosselin et al., 2017). Additionally, RNA sequencing data from iPSC-MG cells and human tissue did not show major overall differences in homeostatic, interferon, activated or NF-κB response genes although varied gene expression was observed among patients (Figure S3A-S3H). These data suggest that the *C9orf72* HRE does not alter the inherent microglial-enriched transcriptional profile indicating that *C9orf72* ALS/FTD and control iPSC-MG share a similar cell-type specific transcriptome in the absence of surrounding CNS cell types, including neurons and macroglia, such as astrocytes and oligodendrocytes.

**Figure 2.**
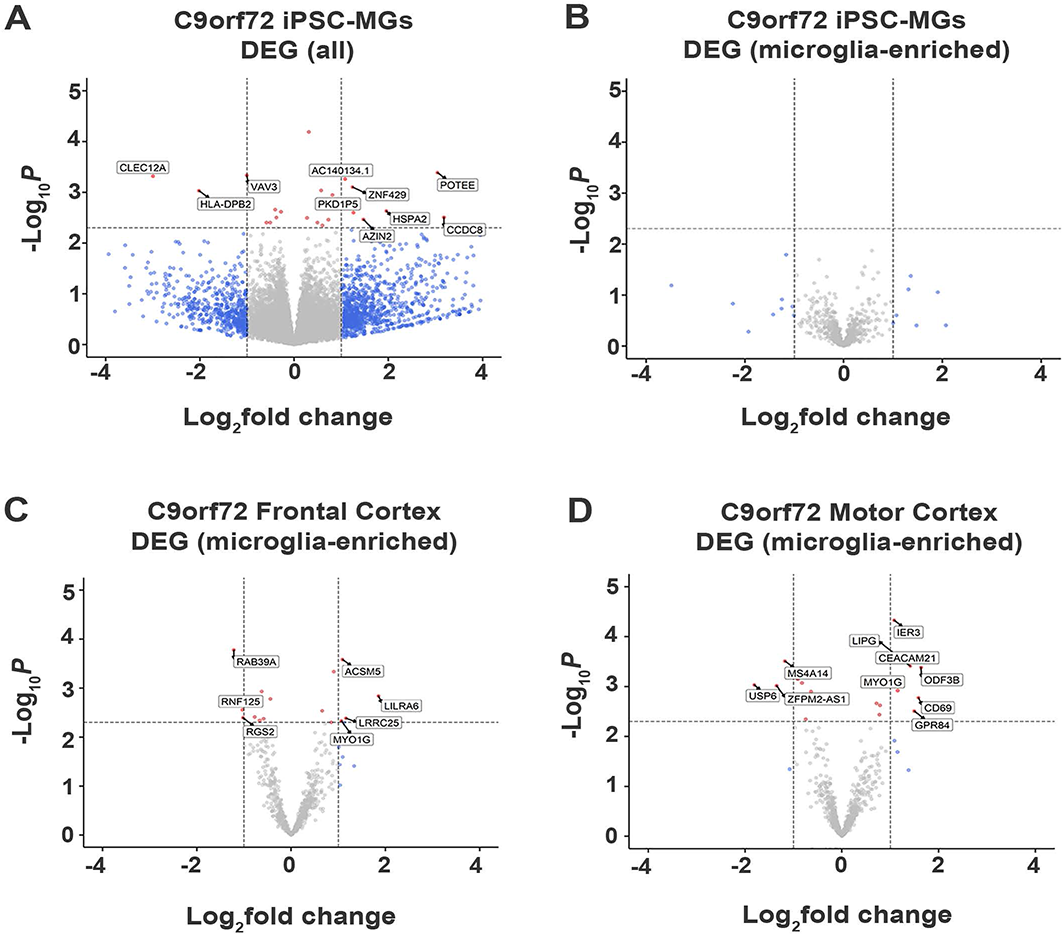
RNA sequencing analyses revealed minor transcriptional changes in mono-cultures of *C9orf72* ALS/FTD iPSC-MG and in postmortem brain tissues of *C9orf72* ALS/FTD patients. Illumina RNA sequencing analysis on mature (DIV 40) iPSC-MG from both, *C9orf72* ALS/FTD patient and healthy controls were compared for overall gene expression changes and for the presence of dysregulated microglial-enriched genes, as defined recently (Gosselin et al., 2017). Additionally, human motor cortex and frontal cortex RNA sequencing data sets obtained through Target ALS and the New York Genome Center were used to identify transcriptional changes. (A) Volcano plot showing differentially expressed transcripts between healthy control (n=4) and *C9orf72* ALS/FTD (n=7) from the full iPSC-MG transcriptome (unadjusted p value <0.005; log_2_ fold change (FC) ± 1). Using these selection criteria, 20 genes were found to be differentially expressed in *C9orf72* ALS/FTD iPSC-MG. (B) Volcano plot of differentially expressed microglial-enriched transcripts (total of 881) indicates that there are no significant expression changes of these particular genes in mono-cultures of *C9orf72* ALS/FTD iPSC-MG (n=7) compared to healthy control (n=4) (unadjusted p value <0.005; log_2_ fold change (FC) ± 1). (C-D) Existing RNA sequencing data from postmortem autopsy brain tissues (frontal cortex, motor cortex of *C9orf72* ALS/FTD patients and controls (Target ALS data set) were also analyzed for differentially expressed 881 microglial-enriched transcripts. Few differentially expressed microglia transcripts in *C9orf72* ALS/FTD were observed compared to controls (seven genes for frontal cortex and 10 genes for motor cortex). Number of brain tissue samples included in the analyses: frontal cortex (control n=16; *C9orf72* ALS/FTD n=8), motor cortex (control n=15; *C9orf72* ALS/FTD n=12) and occipital cortex (control n=4; *C9orf72* ALS/FTD n=5) (unadjusted p value <0.005; log_2_ fold change (FC) ± 1). Volcano plots were generated from DESeq2 output using EnhancedVolcano. All statistical analysis was done in R (version 3.6.2) using raw counts matrices from featureCounts as input. Low expression genes were filtered such that genes with mean read counts < 10 were removed from the analysis. Differential expression analysis was done using DESeq2 (version 1.26.0) using disease status as the model. Tissue data from Target ALS was downloaded from the New York Genome Center (https://www.nygenome.org/) as raw fastq files and pushed through an identical analysis pipeline as data generated for the iPSC-MGs.

To support this hypothesis, we analyzed existing RNA sequencing datasets obtained through the Target ALS consortium and the New York Genome Center. We evaluated gene expression changes of the 881 microglial-enriched RNA transcripts in frontal cortex and motor cortex brain tissues of *C9orf72* ALS/FTD patients (Figures 2C and 2D; Figure S2B; Table S4). We also quantified gene expression changes in the occipital cortex as a brain region considered to be disease-unaffected in ALS/FTD (Figures S2C and S2D; Table S4). Notably, of the 881 microglial-enriched genes, seven were dysregulated in frontal cortex: *LILRA6*, *LRRCA*, *MYO1G*, *RAB39A, ACSM5, RNF125* and *RGS2*. Ten microglial-enriched genes were dysregulated in motor cortex (e.g., *CD69*, *CEACAM21*, *GPR84*, *LIPG* and *MYO1G)* and seven in the occipital cortex (e.g., *CD69*, *EGR2*, *GPR183*, *RGS1* and *SPP1*; Figures 2C and 2D; Figures S2B, S2C and S2D).

### *C9orf72* ALS/FTD iPSC-MG have reduced C9orf72 protein levels, express *C9orf72* hexanucleotide repeat expansion-associated RNA foci and poly-(GP) DPR protein

Glial cells have been shown to exhibit *C9orf72* associated phenotypes in postmortem tissue, including HRE-associated RNA foci and DPR proteins, albeit to a lesser extent than neurons (Sareen et al., 2013, Mizielinska et al., 2013, Lagier-Tourenne et al., 2013b, Mackenzie et al., 2013, Ash et al., 2013, Zhao et al., 2020). To evaluate *C9orf72* iPSC-MG for *C9orf72*-specific disease mechanisms, we first tested *C9orf72* iPSC-MGs for haploinsufficiency by examining the levels of the *C9orf72* transcript in *C9orf72* ALS/FTD and control iPSC-MG. Many studies have reported *C9orf72* haploinsufficiency in patient tissue, however variable results have been observed in human patient-derived *C9orf72* iPSC-differentiated cells, including neurons and astrocytes (Sareen et al., 2013, Donnelly et al., 2013, Zhao et al., 2020). No significant differences in *C9orf72* transcript levels were detected between control and *C9orf72* groups in our RNA seq dataset (Figure 3A) or by quantitative RT-PCR of *C9orf72* (Figure 3B), which is consistent with previous published data (Sareen et al., 2013, Zhao et al., 2020). To measure C9orf72 protein levels we performed quantitative western blot analysis from control and *C9orf72* ALS/FTD iPSC-MG lysates, which revealed a significant reduction in C9orf72 protein in *C9orf72* ALS/FTD iPSC-MGs and was rescued in an isogenic control line (Figures 3C and 3D).

**Figure 3.**
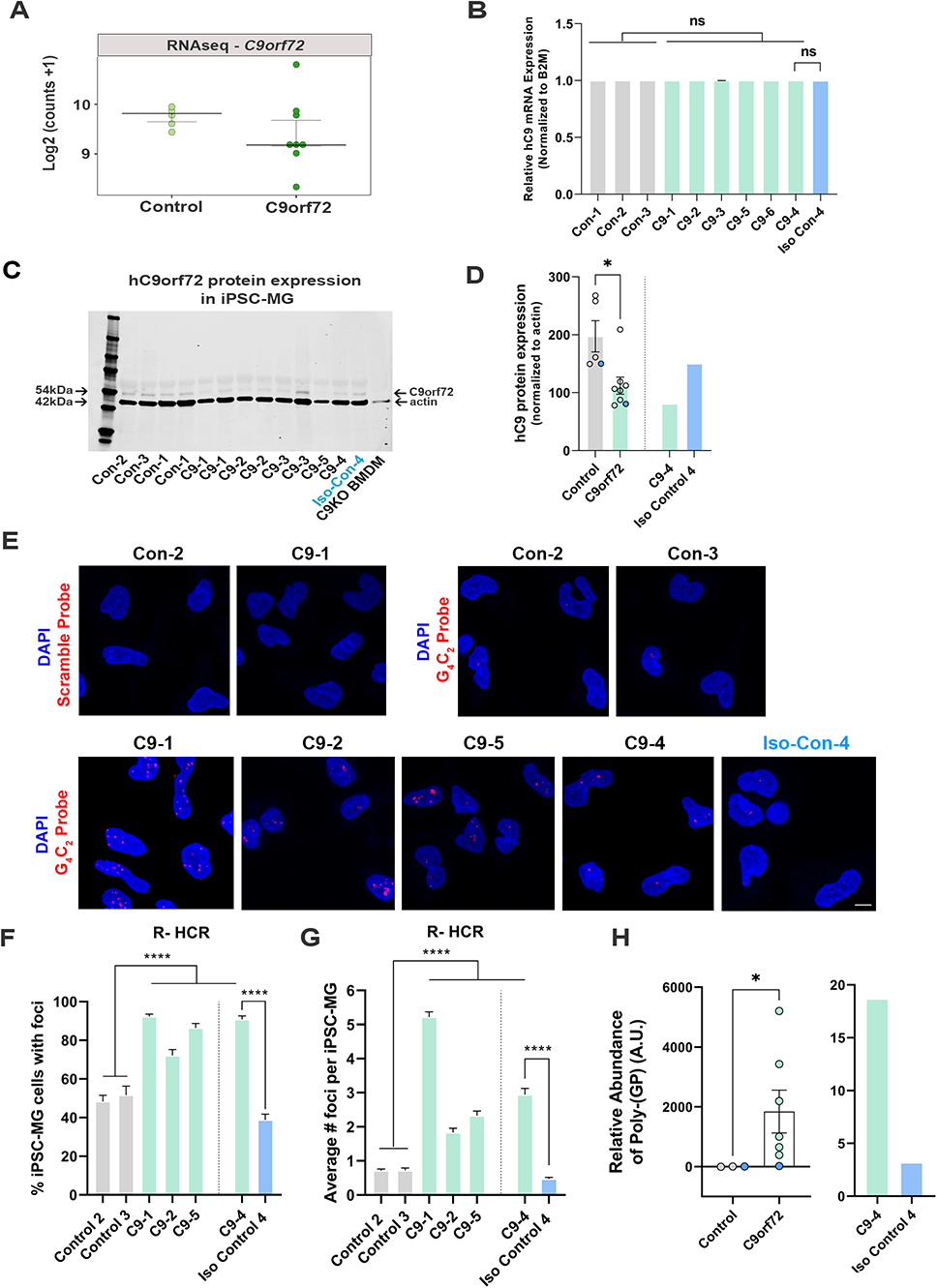
*C9orf72* ALS/FTD iPSC-MG exhibit reduced C9orf72 protein expression, present intranuclear HRE-associated RNA foci and produce poly-(GP) DPR protein. RNA sequencing analysis, qRT-PCR, repeat-hybridization chain reaction (R-HCR), Western Blots (WB) were performed to determine changes in *C9orf72* gene, presence of repeat associated RNA foci and C9 protein expression levels. In addition, an ELISA assay was used to measure poly-(GP) abundance in iPSC-MG. (A) Dot plot showing *C9orf72* level of expression as log2 (counts +1) in control and *C9orf72* ALS/FTD iPSC-MG (Control, n=4 lines; *C9orf72*, n=7 lines; multiple differentiations per line are shown as individual data points; p-value = 0.66, Student’s t-test). No significant differences were observed. (B) Relative human *C9orf72* mRNA expression in control and *C9orf72* ALS/FTD iPSC-MG (normalized expression to beta-2-microglobulin (B2M) in control, n=3 lines, n=1-4 differentiations per line; *C9orf72*, n=6 lines, n=1-3 differentiations per line). No significant differences between groups were detected, including an isogenic control paired with its parent line (C9-4/Iso Con-4). (C) Western blot analysis shows a reduction in human C9orf72 protein expression in *C9orf72* ALS/FTD iPSC-MG. 54kDa C9orf72 protein band and 42kDa actin protein band used as loading control (control, n=4 lines, including an isogenic control-4, n=1-2 differentiation per line; *C9orf72*, n=5 lines, n=1-2 differentiation per line). Bone marrow derived macrophages (BMDM) from a *C9orf72* knockout mouse was used to validate the antibody used for the western blot analysis. (D) Quantification of C9orf72 western blot analysis revealed significant reduction in C9orf72 protein levels (C9orf72 protein expression normalized to actin in control, 197.4, n=4 lines, n=1-2 differentiation per line; *C9orf72*, 112.1, n=5 lines, n=1-2 differentiation per line; p-value = 0.0115; Student’s t-test). Note the rescue of C9orf72 protein levels in the isogenic control line Iso Con-4 vs C9-4 (depicted in blue). (E) R-HCR was performed on iPSC-MG. Representative images of control and *C9orf72* iPSC-MG treated with scramble or *C9orf72*-(G_4_C_2_)_6_ initiator probes. Images show the presence of repeat associated RNA foci in C9 iPSC-MG. *C9orf72*-(G_4_C_2_)_6_ probe hybridized to the corresponding G_4_C_2_ repeats, while absence of RNA foci was observed for the scramble probe. Scale bar = 20µm. (F) Quantification of the percentage iPSC-MG with detectable G_4_C_2_ RNA foci. A significant difference was observed for all C9 iPSC-MG lines when compared to control lines, including C9-4 as compared with corrected isogenic line 4 (control, n=3 lines, including an isogenic control-4; *C9orf72*, n=4 lines, One way ANOVA with multiple comparisons and Student’s t-test were perfomed, p<0.0001). (G) Quantification of RNA foci number per iPSC-MG cell line. Evident increase in RNA foci number in C9 iPSG-MG when compared to control lines (control, n=3 lines, including an isogenic control-4; *C9orf72*, n=4 lines, One way ANOVA with multiple comparisons and Student’s t-test, p<0.0001). (H) A significant increase in intracellular levels of poly-(GP) was detected in *C9orf72* ALS/FTD iPSC-MG using an ELISA assay (relative abundance of poly-(GP) in control, 5.46, n = 3 lines; *C9orf72,* 46.18, n = 7 lines, p=0.0167, two tailed Mann Whitney test). Data is presented as Median ± SEM for *C9orf72* level expression. Student’s t-test was performed for qRT-PCR, WB analysis and RNA foci between C9-4/Iso Control-4. Data is presented as Mean ± SEM. One way ANOVA followed by Tukey’s multiple comparison test was performed for RNA foci analysis, p<0.0001. Two tailed Mann Whitney test statistical analysis was done for the poly-(GP) ELISA assay. \*\**p*-value < 0.01, * *p*-value < 0.05. Exact p-values are reported in the Fig legend.

We then performed repeat-hybridization chain reaction (R-HCR) in mature *C9orf72* iPSC-MG, a more sensitive approach to detect the presence of intranuclear RNA foci (Glineburg et al., 2021) (Figure 3E-G). All *C9orf72* iPSC-MG contained repeat associated intranuclear RNA foci (Figure 3F) with an average number of two to five foci per cell for *C9orf72* iPSC-MG and an average of less than one foci per cell for control iPSC-MG (Figure 3G). A notable and significant difference was observed for C9-4 iPSC-MG compared to its isogenic control (Figure E-G).

Sense and antisense GGGGCC/CCCCGG repeat associated non-AUG (RAN) translation produces five different DPR proteins that accumulate in cells and are proposed to contribute to neuronal toxicity and cellular dysfunction (Zhang et al., 2016b, Freibaum and Taylor, 2017, Gendron et al., 2017, Jiang and Cleveland, 2016, O’Rourke et al., 2015, Zu et al., 2013). Here, we assessed the presence of poly-(GP) in mature iPSC-MG cultures by measuring poly-(GP) abundance in cell lysates using a customized immunoassay (Andrade et al., 2020). We detected a significant increase in poly-(GP) levels in *C9orf72* ALS/FTD iPSC-MGs compared to controls, including the isogenic pair C9-4 (Figure 3H), showing for the first time that the *C9orf72* HRE translates into DPR proteins in endogenous *C9orf72* HRE expressing microglial mono-cultures.

Cytoplasmic TDP-43 inclusions are the hallmark pathology of ALS and FTD and have been reported in glial cells of *C9orf72* ALS/FTD postmortem tissues (Cooper-Knock et al., 2012, Al-Sarraj et al., 2011, Schipper et al., 2016). To evaluate whether iPSC-MG mono-cultures exhibit cytoplasmic TDP-43 inclusions, we performed immunocytochemistry for TDP-43 and measured the nucleocytoplasmic (N/C) ratio of TDP-43 using confocal microscopy. No significant difference in the TDP-43 N/C ratio was observed between control and *C9orf72* ALS/FTD microglia at 40 DIV (Figures S4A and S4C). One explanation for TDP-43 mislocalization is a defect in nucleocytoplasmic trafficking leading to the accumulation of TDP-43 protein in the cytoplasm (Moore et al., 2020, Zhang et al., 2015, Chou et al., 2018). Our laboratory has recently shown that another RNA binding protein, the RNA editing enzyme adenosine deaminase acting on double stranded RNA 2 (ADAR2), is mislocalized to and accumulates in the cytoplasm of motor neurons in *C9orf72* ALS/FTD (Moore et al., 2019). We therefore wondered whether *C9orf72* microglia would display similar ADAR2 mislocalization. Like TDP-43, immunocytochemistry of *C9orf72* ALS/FTD iPSC-MG for ADAR2 revealed no nucleocytoplasmic mislocalization resulting in an unchanged N/C ratio in microglia mono-cultures (Figures S4B and S4D). These data suggest that while *C9orf72* ALS/FTD iPSC-MG mono-cultures do exhibit *C9orf72* pathobiological phenotypes as shown in *C9orf72* ALS/FTD iPSC neurons, there are no microglial cytoplasmic inclusions of TDP-43 or ADAR2.

### IPSC-MG carrying *C9orf72* HRE do not alter their inflammatory response to LPS stimulation

Microglia are able to exacerbate or promote neurodegeneration by releasing pro-inflammatory cytokines, including interleukin-1 (IL1), interleukin-6 (IL6) and tumor necrosis factor-α (TNFα), or anti-inflammatory cytokines such as IL-4 and IL-10 (Smith et al., 2012, Moreno-Martinez et al., 2019, Olesen et al., 2020). As a result, neuroinflammation is considered a major contributor to neuronal dysfunction in neurodegenerative diseases, including ALS/FTD (Olesen et al., 2020, McCauley and Baloh, 2019, Beers and Appel, 2019, Lall and Baloh, 2017). To investigate if *C9orf72* ALS/FTD iPSC-MG respond differently to extracellular stimuli, we treated iPSC-MGs with LPS, an endotoxin used to evoke immune responses *in vitro* and *in vivo* (Abud et al., 2017, Hong et al., 2020, Beurel and Jope, 2009, Furube et al., 2018). Following LPS treatment, we tested for the presence of released chemokines and cytokines in the iPSC-MG cell culture supernatants, as described recently (Abud et al., 2017). Upon LPS stimulation, both control and *C9orf72* ALS/FTD iPSC-MG mono-cultures exhibited increased release of IL1α, IL1β, IL6 and TNFα compared to basal conditions but no differences between the experimental groups were observed (Figure 4A-4D; individual cell response shown in Figure S5A-S5D). These results confirm the ability of *C9orf72* ALS/FTD patient-derived iPSC-MG mono-cultures to respond to extracellular stimuli and support previous *in vitro* studies showing no significant changes in inflammatory responses in murine microglia cell lines overexpressing *C9orf72* HRE (Rostalski et al., 2020). Additional sets of iPSC-MGs were treated with LPS and analyzed for gene expression alterations by RNA sequencing. Both, healthy control and *C9orf72* microglia responded to LPS treatment with significant gene expression changes (Figure S6A-S6B; log_2_ fold change (FC) ± 1, p value <0.05). Expected dysregulations were found in genes involved in inflammatory processes and interferon gamma signaling pathways, e.g. *CXCL8*, *CXCL10*, *TNIP3*, *CCL4L2*, *CCL20*, *IFIT2*, *IFIT3*, *IL23A*, *IL12B* and *IL6*. Among the top significantly dysregulated disease-specific genes is the cytokine IL-1α, which is highly expressed and secreted by microglia and is known to be a strong inducer of an A1 reactive astrocyte phenotype (Figure 4E; Figure S5E; log_2_ fold change (FC) ± 1, p value <0.05) (Zhang et al., 2014, Bennett et al., 2016, Liddelow et al., 2017).

**Figure 4.**
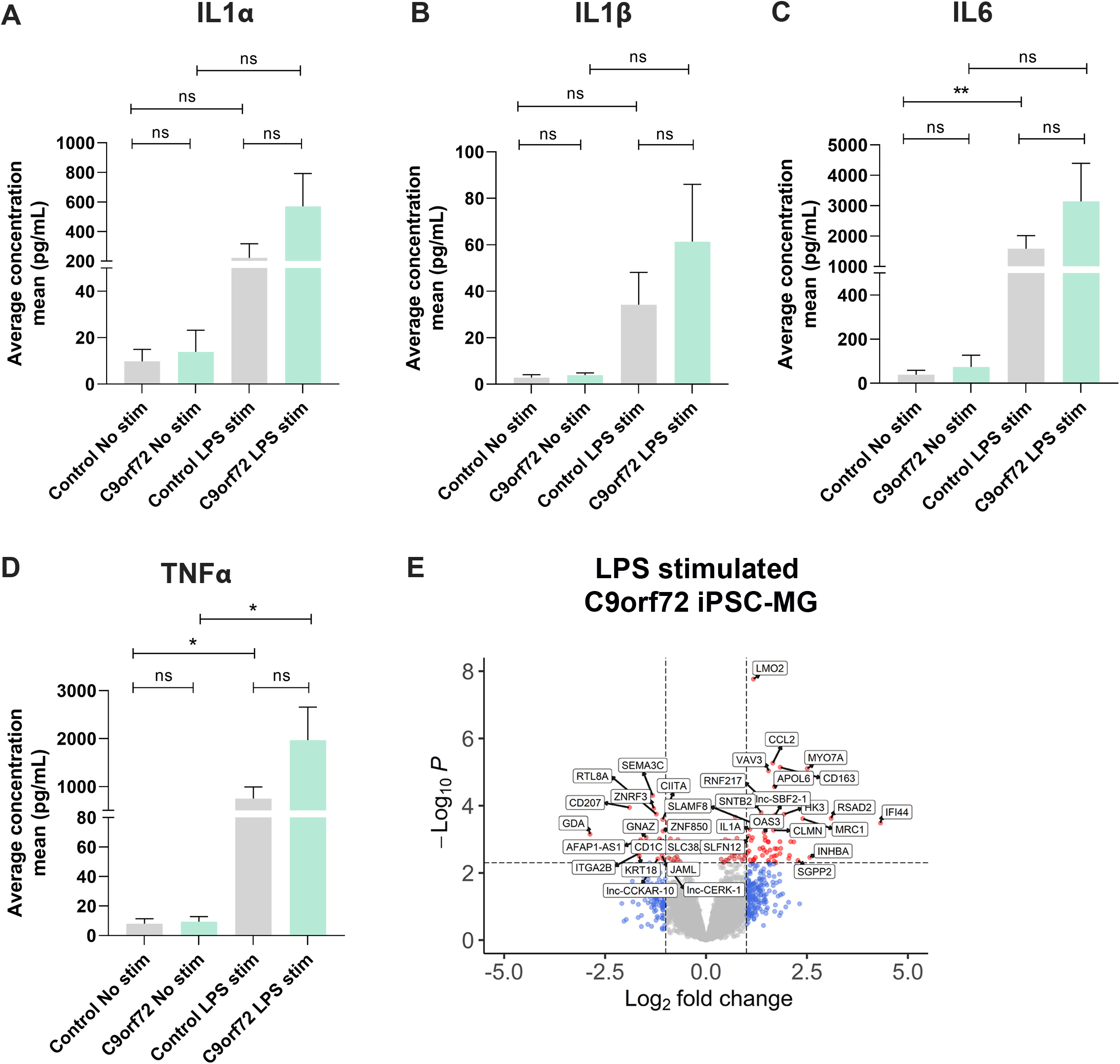
*C9orf72* ALS/FTD and healthy control iPSC-MG show equal response to LPS stimulation. *C9orf72* ALS/FTD iPSC-MG and control iPSC-MGs were treated with LPS (100ng/ml) for 6hr. The conditioned media was collected and analyzed for a cytokine/chemokine profile using the U-plex Biomarker Group 1 (Human, Mesoscale) Multiplex Assay. (A-D) Under basal conditions, *C9orf72* ALS/FTD iPSC-MG monocultures respond with an increase in IL-1α, IL-6 and TNF-α inflammatory cytokines similar to controls (See Figure S5A-S5D for response per replicate within a sample). Data presented as the average concentration of secreted proteins (pg/ml), Mean ± SEM, n=4 control lines, 2-3 replicates; n=3 *C9orf72* lines, 3 replicates; exact *p*-values displayed in the graphs; \**p* ≤ 0.05; \*\**p* ≤ 0.01; \*\*\*\**p* ≤ 0.0001; Student’s t-test. (E) Volcano plot showing differentially expressed genes in *C9orf72* ALS/FTD (n=4) from 6 h LPS stimulation when compared to healthy control (n=5) (See Figure S5E). Volcano plot was generated with same criteria as aforementioned.

### *C9orf72* ALS/FTD iPSC-MGs exhibit normal phagocytic activity

A major function of microglia is to clear unwanted toxic substances and cell debris that can negatively impact brain function via phagocytosis. To determine whether control and *C9orf72* ALS/FTD iPSC-MG differ in their phagocytic activity and ability to clear toxic products we measured iPSC-MG engulfment of a fluorescently tagged synthetic amyloid beta (Aβ) cargo, which has been shown previously to be phagocytosed by microglia (Paresce et al., 1996, Paolicelli et al., 2017). We transiently treated iPSC-MGs for 5 minutes with 1µM Aβ (1-40)-TAMRA or vehicle and then assessed engulfment and clearance of Aβ (1-40)-TAMRA at different time points after Aβ washout (0, 30, 60 min) using fluorescent confocal microscopy. To quantify this engulfment, we performed immunohistochemistry on iPSC-MG for TREM2, an immune receptor selectively expressed in microglia (Figure 5A-5F). We calculated the percentage of iPSC-MG surface area covered by Aβ (1-40)-TAMRA protein in individual cells at the 5 min and 1 h time points. We observed varying levels of retention of phagocytosed Aβ depending on the patient lines at both time points, with four lines out of seven lines showing significant differences compared to healthy control lines. While variable, patient lines showing aberrant phagocytic Aβ uptake also showed loss of C9orf72 protein (see Figures 3C and 3D). Altered Aβ uptake dynamics suggest potential dysfunction of the endosomal-lysosomal pathways and inability to adequately process phagocytosed cargo in select patient lines (Figures 5H and 5J). When grouped together, no differences in phagocytosed content between *C9orf72* and healthy control iPSC-MG was noted (Figures 5I and 5K). To take into account possible changes in cell size, we measured total microglia cell surface. No significant differences in the surface area between experimental groups were observed (Figure 5G). Finally, we wondered whether microglia would respond to toxic protein exposure with an altered transcriptional profile. Therefore, iPSC-MGs were treated with Aβ (1-40)-TAMRA and analyzed for gene expression changes via RNA sequencing analyses. Even with lower stringency for DEG analyses, neither control nor disease iPSC-MG showed large numbers of dysregulated genes when we compared treated to untreated cells within each experimental group (Figure S6C and S6D; log_2_ fold change (FC) ± 1, p value <0.05). However, when we examined DEG in Aβ-treated *C9orf72* iPSC-MG in reference to Aβ-treated control microglia, we noted several significantly differentially expressed genes (41 genes; log_2_ fold change (FC) ± 1, p value <0.05; Figure 5L; Figure S6E). Among the upregulated genes were *FCGR3A* and *DOCK1*, genes known to play a role in phagocytosis (Sivagnanam et al., 2010, Wu and Horvitz, 1998). Additional upregulated genes were found to play a role in endocytic trafficking, cytoskeletal rearrangement, or cell migration (*RAB15*, *AHNAK* and *SDC3*) (Zuk and Elferink, 2000, Dumitru et al., 2013, Carey, 1996).

**Figure 5.**
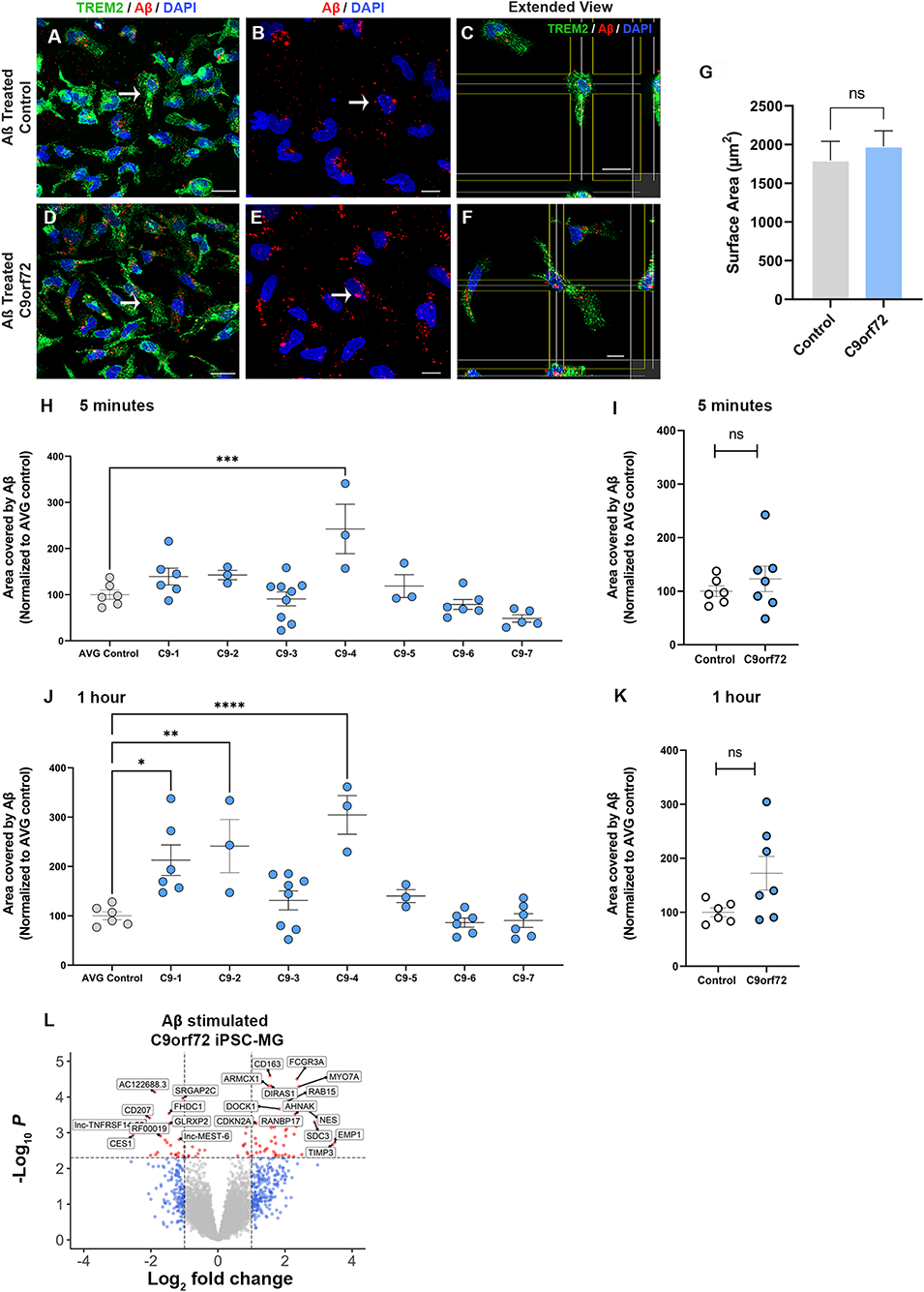
*C9orf72* ALS/FTD iPSC-MGs show increased phagocytosis of synthetic Aβ (1-40) in select patient lines. Control and *C9orf72* ALS/FTD iPSC-MG were exposed to 1 μM Aβ (1-40) TAMRA for 5 min, washed and analyzed for phagocytosed content at 5 min and 1 h. Cells were immunostained for microglia marker TREM2 and images were taken using confocal microscopy. Imaris imaging software was used to illustrate and quantify the engulfed protein. (A, D) Representative images of healthy controls and *C9orf72* ALS/FTD iPSC-MG stained for TREM2 (green) and highlighting Aβ (1-40) TAMRA (red) inside the cells at 30 minutes. White arrow point at phagocytic cells containing Aβ (1-40) TAMRA (red). Scale bar, 20μm. (B, E) Aβ (1-40) TAMRA (red) internalized by healthy controls and *C9orf72* ALS/FTD iPSC-MG. Scale bar, 20μm. (C, F) Extended view showing phagocytic activity in both iPSC-MG groups. Scale bar, 20μm. (G) No significant differences in the microglia cell surface area were observed upon Aβ exposure (control, n=5 lines; *C9orf72*, n=6 lines; n= 1-2 differentiations per line; 2-3 replicates; 6-7 pictures per replicate; n= 6-10 cells per picture; p-value = 0.58 using Student’s t-test). (H-K) Select patient lines showed significant differences in the percentage of cell surface area covered by Aβ (1-40)-TAMRA after 5 minutes (1 out of 7 lines) or 1 h (3 out of 7 lines) (control, n=6 lines; *C9orf72*, n=7 lines; n= 1-2 differentiations per line; 2-3 replicates; 6-7 images per replicate; n= 6-10 cells per image). One-way ANOVA followed by a Tukey’s multiple comparison test was performed, \**p* ≤ 0.05; \*\**p* ≤ 0.01; *** *p* ≤ 0.001; \*\*\*\**p* ≤ 0.0001. (L) Differentially expressed genes from *C9orf72* ALS/FTD Aβ (1-40) treated iPSC-MG (control, n=5 lines; *C9orf72*, n=4 lines; see Figure S6E). Volcano plots were generated from DESeq2 output using EnhancedVolcano. All statistical analysis was done in R (version 3.6.2).

### *C9orf72* ALS/FTD iPSC-MGs engulf human healthy brain synaptoneurosomes similar to control iPSC-MGs

An important aspect of microglia-neuron communication in neurodegeneration is the role of microglia in the maintenance and refinement of synaptic networks through the selective pruning of synapses. This process occurs predominantly during development (Stevens et al., 2007, Bialas and Stevens, 2013, Tremblay et al., 2011). However, synaptic pruning pathways are known to be re-activated in neurodegeneration leading to synapse loss and contributing to cognitive impairments (Hong et al., 2016, Lui et al., 2016, Colom-Cadena et al., 2020, Henstridge et al., 2016). To determine if iPSC-MG can phagocytose synapses and to test whether *C9orf72* microglia exhibit altered phagocytosis due to the *C9orf72* HRE, we exposed iPSC-MG mono-cultures to healthy control human brain synaptoneurosomes (hSN) and followed synaptoneurosomes engulfment via live confocal microscopy. Fresh frozen control human postmortem tissues were used to prepare hSN. Synaptic fractions were enriched for pre-synaptic protein synaptophysin and the post-synaptic density 95 (PSD-95) compared to total brain homogenate and nuclear marker histone 3 (Figures S7A and S7B). Control and *C9orf72* ALS/FTD iPSC-MGs were labeled with the live cell nuclear marker Hoechst (Figure 6A-6I) to identify individual cells followed by treatment with hSN fluorescently tagged with pHrodo succinimidyl ester (hSN-rodo; Figure 6A; Figures 6J-6Q). An increase in pHrodo fluorescence is indicative of an uptake of hSN-rodo into acidic intracellular compartments of iPSC-MGs. An initial six hour live cell imaging determined a distinct rapid internalization of hSN-rodo in control and *C9orf72* ALS/FTD iPSC-MGs with more than 60% of cells exhibiting phagocytic activity at two hours (Figure 6R). iPSC-MG phagocytosis was reduced significantly in the presence of cytochalasin-D, an actin polymerization inhibitor known to inhibit phagocytosis (Figures 6R and 6S). We then treated additional sets of iPSC-MGs with hSN-rodo and performed fluorescent live cell imaging for two hours, capturing images every 10 min (Figure S7C and Movie S1). The mean intensity of hSN-rodo was similarly increased in *C9orf72* ALS/FTD iPSC-MGs compared to controls (Figures 6T and 6U). Together, these data suggest that *C9orf72* ALS/FTD iPSC-MGs exhibit known microglia phagocytic activity as shown by the engulfment of healthy human brain synaptoneurosomes, similar to what we showed for Aβ above. Additional studies are required to determine if *C9orf72* ALS/FTD iPSC-MGs respond differently to diseased human brain synaptoneurosomes, which may contain specific signaling molecules necessary for microglia activation and elimination of synapses.

**Figure 6.**
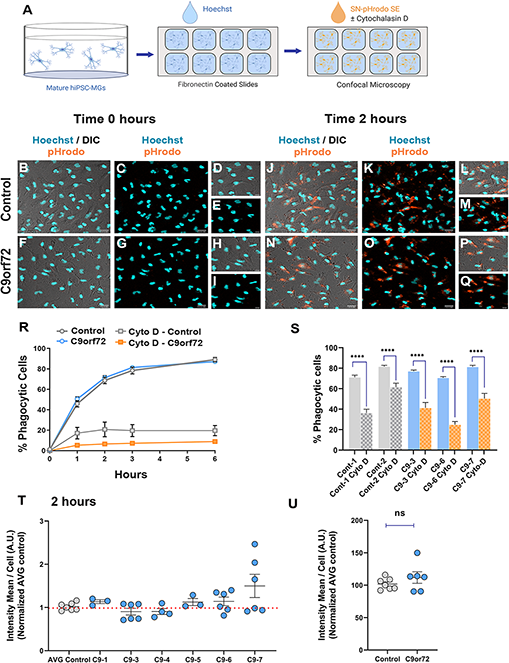
Phagocytic uptake of human brain synaptoneurosomes by iPSC-MGs. Live cell imaging of control and *C9orf72* ALS/FTD iPSC-MGs engulfing pHrodo-labeled human brain synaptoneurosomes was performed over a 2 h time period with pictures taken every 10 minutes. (A) Illustration of iPSC-MGs treated with human brain synaptoneurosomes. Cultured control and *C9orf72* ALS/FTD iPSC-MGs (Day 40) are plated onto fibronectin coated slides and labeled with Hoechst nuclear marker followed by hSN-rodo treatment ± 10μM Cytochalasin D (actin polymerization inhibitor). (B-C) Control and (F-G) *C9orf72* ALS/FTD iPSC-MG images taken at 0 h time point. IPSC-MGs labeled with live nuclear stain Hoechst (cyan) and exposed to hSN-rodo (orange). Differential interference images (DIC) enhance the visualization of individual iPSC-MG morphology (B, F). No evident phagocytosis of hSN-rodo is seen at this time point (C, G). Higher magnification representative images at 0 h time point of control (D-E) and (H-I) *C9orf72* ALS/FTD iPSC-MGs. Scale bars = 50μm. (J-K) Representative images of control and (N-O) *C9orf72* ALS/FTD iPSC-MG at 2 h show an increase in hSN-rodo fluorescent signal inside iPSC-MG indicative of phagocytosis and uptake into intracellular acidic compartments. Scale bar = 40µm. (L-M; P-Q) Higher magnification representative images highlighting the increase in hSN-rodo signal in individual iPSC-MG at 2 h. Scale bar = 10µm. (R) Percentage of iPSC-MG engulfing hSN-rodo during the 6 h initial time course of live imaging. At 2 h, control and *C9orf72* iPSC-MG showed 69% and 71% phagocytic activity, respectively. No significant differences in synaptoneurosome uptake was observed between groups. 10uM cytochalasin D, an actin polymerization inhibitor was used as a negative control to inhibit phagocytic activity in iPSC-MGs (For 6 h initial time course; control, n=1 line; *C9orf72,* n=2 lines; 3 replicates; n =6 images/ replicate/ time point; n=8 cells per image for hSN-rodo; for cytochalasin D, n =6 images/group/ time point n = 10 cells per image). (S) Percentage of iPSC-MG engulfing hSN-rodo at 2 h time point with cytochalasin D treatment. A significant decreased in phagocytic activity was observed in controls and *C9orf72* ALS/FTD iPSC-MG in the presence of cytochalasin D (Cyto-D, control, n=2 lines; *C9orf72,* n=3 lines, n =6 images/group; \*\*\*\**p* ≤ 0.0001 using Student’s t-test). (T-U) HSN-rodo mean intensity per cell normalized to average control at 2 h (control, n=7 lines; *C9orf72*, n=6 lines; 3 replicates; 6-7 images per replicate; n= 6-10 cells per image). Each iPSC-MG line was normalized and compared to the average controls. Using Student’s t-test, we observed no differences in the phagocytosis of control human brain synaptoneurosomes, Data presented as Mean intensity ± SEM.

## DISCUSSION

An extensive body of evidence has suggested that glia contribute to the neurodegeneration observed in ALS and FTD (Ilieva et al., 2009, Yamanaka et al., 2008, Ban et al., 2019, Filipi et al., 2020, Valori et al., 2019, Dols-Icardo et al., 2020). Transcriptional assessments and proteomic approaches across the ALS/FTD spectrum using predominantly postmortem autopsy tissue have reported robust glia signatures and glia protein modules, respectively, emphasizing a glial involvement in inflammation and contribution to disease (Umoh et al., 2018, Tam et al., 2019, D’Erchia et al., 2017, Dols-Icardo et al., 2020).

In the present study, we generated microglia from *C9orf72* ALS/FTD patient-derived iPSC to evaluate their cellular and molecular phenotypes. The differentiation protocol was selected based on the transcriptional and functional similarities between the generated human iPSC-MG to adult human microglia as well as for their high purity, yield and distinction from other myeloid cells such as monocytes and dendritic cells (McQuade et al., 2018, Abud et al., 2017). The *C9orf72* ALS/FTD iPSC-MG displayed classic microglia characteristics and presented a unique transcriptomic signature profile compared to iPSC-CNs or other glial cell types (Zhang et al., 2014). Applying a list of human microglia-enriched genes from Gosselin and colleagues (Gosselin et al., 2017), transcriptional analyses revealed no significant differences of these microglial genes between *C9orf72* ALS/FTD and control iPSC-MG under basal, unstimulated culture conditions. This supports the notion that the presence of the *C9orf72* HRE does not affect iPSC microglia differentiation and their baseline transcriptome.

As microglia function is influenced by the cellular environment, we examined changes of previously reported microglial-enriched genes (Gosselin et al., 2017) using existing bulk RNA sequencing data from postmortem *C9orf72* ALS/FTD brain tissues. Surprisingly, no significant changes in the microglial-enriched genes were found either in the frontal, motor or occipital cortex supporting the need for single-cell resolution technologies to better identify changes in gene expression in specific cell populations, similar to what has been reported in Alzheimer’s disease brain tissues’ analyses (Keren-Shaul et al., 2017, Mrdjen et al., 2018, Masuda et al., 2020, Bottcher et al., 2019, Masuda et al., 2019, Sankowski et al., 2019). Cell type-specific analyses from postmortem *C9orf72* ALS/FTD brain tissue will further allow for the identification of subsets of microglial subpopulations and associate their transcriptional signatures with potential neuroprotective or detrimental roles, as well as microglial-specific disease pathways and mechanisms.

To our knowledge, this data is the first to indicate that human endogenous *C9orf72* ALS/FTD iPSC-MG exhibit intrinsic *C9orf72* pathology. Although, transcriptional analyses indicated variability of *C9orf72* mRNA levels across *C9orf72* ALS/FTD iPSC-MG patient lines, no significant differences in *C9orf72* expression were observed by RNA sequencing or quantitative RT-PCR analysis. Our data is consistent with previous studies in iPSC patient-derived neurons and astrocytes (Sareen et al., 2013, Zhao et al., 2020). A larger sample size might be required for RNA sequencing analysis to obtain significant results. As for the qRT-PCR, due to the existence of several *C9orf72* RNA variants and the notion that there is cell type-specific promoter usage of *C9orf72* variants, the primers used for the present qRT-PCR might not reflect a microglia-specific reduction in *C9orf72* gene expression (Sareen et al., 2013). We observed a significant decrease in C9orf72 protein expression in *C9orf72* iPSC-MG, which is an interesting finding, as previous reports on *C9orf72* iPSC-astrocytes showed no reduction in C9orf72 protein levels (Zhao et al., 2020). The loss of function of C9orf72 protein has been implicated in alterations of endosomal-lysosomal pathways, hence could contribute to the phagocytic differences we observed amongst individual *C9orf72* patient lines (Amick et al., 2016, Amick et al., 2020, Sullivan et al., 2016, O’Rourke et al., 2016, Farg et al., 2014, Shi et al., 2018). In support of this hypothesis, patient lines showing significant changes in Aβ phagocytosis had lower C9 protein levels when compared to healthy controls (Figure 3C and 3D; Figure 5J). Additional studies reported enlarged lysosomes in *C9orf72* postmortem patient tissues and proposed that the function of the C9orf72 protein is linked to the regulation of lysosomes and autophagy (O’Rourke et al., 2016, Shao et al., 2019, Amick and Ferguson, 2017, Sullivan et al., 2016, Wang et al., 2020).

We also evaluated *C9orf72* iPSC-MG for the non-canonical translation of DPR proteins, specifically poly-(GP), and are the first to report endogenous poly-(GP) production in patient *C9orf72* microglia, suggesting that similar to *C9orf72* iPSC-astrocytes, microglia undergo repeat-associated non-ATG translation (Zhao et al., 2020). Recent studies have shown neuron-astroglia transmission of *C9orf72* associated DPRs via exosomes in an *in vitro* culture system (Westergard et al., 2016). It is yet to be determined if microglia can similarly contribute to transmission of DPRs to neighboring cells.

*C9orf72* DPRs have been suggested to contribute to nucleocytoplasmic trafficking defects present in *C9orf72* ALS/FTD neurons (Moore et al., 2020, Jovicic et al., 2015, Boeynaems et al., 2016). One of the consequences of these trafficking defects is the mislocalization of nuclear RNA binding proteins, such as TDP-43. TDP-43 pathology has been shown to be present in glia of *C9orf72* postmortem tissues (Cooper-Knock et al., 2012, Schipper et al., 2016, Brettschneider et al., 2014, Fatima et al., 2015, Yamanaka and Komine, 2018). TDP-43 has further been associated with neuroinflammation, microglia neuroprotection and the regulation of microglia phagocytosis (Swarup et al., 2011, Spiller et al., 2018, Paolicelli et al., 2017). In the present study, TDP-43 cytoplasmic accumulations or loss of nuclear TDP-43 was not detected in *C9orf72* ALS/FTD iPSC-MG mono-cultures; similar to what has been observed in *C9orf72* iPSC patient-derived astrocytes (Zhao et al., 2020). Furthermore, we found no significant difference in the nucleocytoplasmic ratio of the RNA editing protein ADAR2, which has recently been shown to be mislocalized to the cytoplasm of *C9orf72* ALS/FTD iPSC motor neurons, as well as in neurons in *C9orf72* ALS/FTD postmortem tissues and *C9orf72* ALS/FTD mouse model brain tissues (Moore et al., 2019). The absence of TDP-43 and ADAR2 mislocalization could be due to cellular age of the differentiated cells, as recent human postmortem tissue studies suggested that TDP-43 mislocalization is a late stage event of ALS pathogenesis (Vatsavayai et al., 2016).

Cerebral spinal fluid and blood cytokine profiles are significantly altered for a large array of cytokines and chemokines in ALS/FTD (Lu et al., 2016). In addition, recent data support a correlation of specific immune responses of gene-associated ALS subgroups to patient survival (Olesen et al., 2020). As for *C9orf72* ALS/FTD, previous studies in *C9orf72* knockout or *C9orf72* deficient mice revealed increased IL-6 and IL1β mRNA levels in microglia and an upregulation of inflammatory pathways, suggesting an association between the loss of *C9orf72* and altered microglia function and pro-inflammatory phenotypes (O’Rourke et al., 2016, Prudencio et al., 2015, Lagier-Tourenne et al., 2013b, Lall et al., 2021). Here, *C9orf72* ALS/FTD iPSC-MG mono-cultures exhibit a comparable response to control iPSC-MG upon LPS stimulation. Further studies are necessary to determine if *C9orf72* ALS/FTD iPSC-MG present or exacerbate an inflammatory phenotype when co-cultured with diseased neurons or astrocytes knowing that a direct contact of iPSC-MG with CNS cells can influence their gene expression (Abud et al., 2017). Likewise, it will be important to determine if an anti-inflammatory response is acquired in the presence of *C9orf72* ALS/FTD iPSC-neurons or other glial cell types.

Overall, we report intrinsic properties of *C9orf72* ALS/FTD iPSC-MG mono-cultures and set the stage for the use of this human endogenous disease model in co-cultures with iPSC neurons and other glial cells, astrocytes and/or oligodendrocytes. *C9orf72* ALS/FTD iPSC-MG can be used in a 2 or 3-dimensional co-culture system, or can be transplanted *in vivo* into mouse models to further assess microglial contribution to neuronal dysfunction and degeneration in *C9orf72* ALS/FTD. Finally, this human cell culture model provides novel opportunities to screen microglial-targeted new therapeutics for future drug development for ALS/FTD patients.

## MATERIAL AND METHODS

### iPSC lines

The majority of the iPSC lines used in our studies were purchased from Cedars Sinai induced pluripotent stem cell core https://www.cedars-sinai.edu/research/areas/biomanufacturing/ipsc.html or obtained from collaborators who have characterized the lines in previous publications. All purchased iPSC line from Cedars Sinai are characterized for: 1) positive staining of pluripotency markers (Oct3/4, NANOG, SOX2, TRA-1-60, TRA-1-81, SSEA4); 2) karyotyping to reveal no chromosomal abnormalities; 3) determine a pluripotency score via PrimeView global gene expression profile assay (PluriTest); 4) trilineage differentiation potential via Taqman hPSC scorecard assay to confirm appropriate expression of ectodermal, endodermal and mesodermal factors in the iPSC cells; 5) cell line authentication analysis - STR Analysis - to confirm identity/profile matching score with primary tissue. A Certificate of Analysis (COA) for these lines is available upon request. Please see Supplemental Table S1 for comprehensive information on the lines used in these studies.

### Generation of Hematopoietic Progenitor Cells (HPCs) DIV -1 to DIV 12

IPSCs were differentiated into microglia following an established protocol (McQuade et al., 2018). Briefly, iPSCs were maintained in mTeSR Plus Kit (Stemcell Technologies # 05825) in 10cm dishes. At DIV -1 of HPCs differentiation, five to seven day old iPSCs cultures were then used to generate cluster of differentiating 43 positive (CD43^+^) hematopoietic progenitor cells (HPCs) following a 12 day commercially available kit (STEMdiff Hematopoietic Kit; Stemcell Technologies # 05310). IPSCs were cleaned and 1/3 dish was gently dissociated with dispase (Stemcell Technologies # 07923) for 12-15 minutes at 37°C. IPSCs were then collected and spun down at 500rpm for 1-2 minutes and resuspended in 2mL of mTeSR Plus media with 20 µM ROCK inhibitor Y-27632 (Stemcell Technologies # 72304). Matrigel, hESC-Qualified Matrix (Corning # 354277) coated six well plates were used to start the HPC differentiation. Using a 5mL serological pipette, one drop or two drops of iPSCs were seeded per well into matrigel. Then, next day, on DIV 0 of HPCs differentiation, wells with 80 small colonies per well were selected to start HPCs differentiation. IPSCs were fed following the manufacturer’s instructions. On DIV 12 of HPCs differentiation, only the non-adherent HPCs were transferred to a new six well plate to start microglia differentiation (DIV 12/0).

### Differentiation of HPCs into microglia cells DIV 12/0 – DIV 40/28

Matrigel, GFR (growth factor reduced) Membrane Matrix (Corning # 356231) coated six well plates were prepared to start microglia differentiation (DIV12/0). HPCs were differentiated into microglia for 28 days using serum free media conditions. HPCs in suspension were collected and spun down at 300 xG for 6 minutes and resuspended into microglia basal media (MBM, 2mL/well) containing: DMEM/F12 no phenol (Gibco # 11-039-021), 2% Insulin Transferin Selenite (Gibco # 41400045), 2% B27 (Gibco # 17504-044), 0.5% N2 (Gibco # 17502-048), 1% Glutamax (Gibco # 35050-061), 1% NEAA (Gibco # 11140-050), 1% Pen/Strep (Gibco # 15140-122), 400μM 1-Thioglycerol (Sigma # M1753), 5μg/mL human insulin (Sigma # I2643) and supplemented with 3 growth factors (GFs): 100ng/mL human recombinant interleukin-34 (IL-34; Peprotech # 200-34), 25ng/mL macrophage colony-stimulating factor (M-CSF; Gibco # PHC9501) and 50ng/mL transforming growth factor β1 (TGFβ1; Miltenyi Biotec # 130-108-969) (MBM + 3GFs); cytokines and growth factors known to be essential for the development of microglia (Butovsky et al., 2014, Greter et al., 2012, Wang et al., 2012, Abud et al., 2017, Pandya et al., 2017, Abutbul et al., 2012, Chen et al., 2002, Elmore et al., 2014). Microglia differentiation starts (DIV 12/0) once the HPCs are transferred and plated at a density of 200 000 cells per well of a six well. Cells will predominantly grow in suspension. On DIV 2, 4, 6, 8 and 10 of microglia differentiation, 1mL of MBM + 3GFs media was added to each well. On DIV 12, a partial media change was done. IPSC-MGs from one six well plate were spun down at 300 xG for 6 minutes and resuspended into MBM + 3 GFs and split back into the same six well plate. On DIV 14, 16, 18, 20, 22, 24, 1mL media was added per well. On DIV 25, MBM + 3GFs was changed to maturation media composed of MBM supplemented with the 5 growth factors (MBM +5 GFs): 100ng/mL IL34, 25ng/mL M-CSF and 50ng/ml TGFβ1, 100ng/mL cluster of differentiation 200 (CD200; Novoprotein # C31150UG) and100ng/mL fracktaline chemokine C-X3-C motif ligand 1 (CX3CL1; Peprotech # 300-31) (MBM + 5 GFs). The presence of CD200 and CX3CL1 in the culture media, both glial and neuronal molecules are critical for microglia maturation and maintenance of an *in vivo-*like microglia resting state phenotype in an *in vitro* setting (Barclay et al., 2002, Kim et al., 2011, Kierdorf and Prinz, 2013) (Figure 1A). At DIV 28, IPSC-MG reached maturation. 1mL of MBM + 5GFs media was added to the cultures every other day. Mature cells were used for experimentation within 10 days (DIV 28-DIV38).

### Differentiation of iPSCs into cortical neurons

For cortical neuron differentiation, 70% confluent iPS cells maintained in mTeSR Plus media on 10 cm dishes were used for embryoid bodies (EBs) formation. The cells were cultured in low attachment six well plates (Greiner bio-one # 657970) using WiCell Medium containing: DMEM/F12 (Gibco #11330057), 25% knock out serum replacement (Gibco # 10828-028), 1.3% L-glutamine (Gibco # 35050-061), 1.3% NEAA (Gibco # 11140-050), 0.1mM 2-Mercaptoethanol (Sigma# M3148) and placed on a shaker in the incubator for 8 days to allow EBs formation. EBs were then resuspended in Forebrain Neural Induction Media (FB-NIM) containing: DMEM/F12, 1% N2 supplement (Gibco # 17502-048), 1% NEAA, 2ug/mL heparin (Sigma # H3149), 10 μg/mL bFGF (Stemcell Technologies # 78003) and plated on to T25 flasks coated with basement membrane matrigel (Corning # 356234) to allow formation of neuronal rosettes. Neuronal rosettes were maintained in FB-NIM for the next 10 days and then, collected and maintained in suspension on a shaker with half FB-NIM media changes every other day to allow for neurosphere growth. Neurospheres were maintained for 20 days in FB-NIM and then resuspended using forebrain neuronal differentiation media (FB-DM) containing: Neurobasal Medium (Corning # 21103-049), 2% B27 (Gibco # 17504-044), 10 ug/mL BDNF (Stem cell technologies # 78005), 10 ug/mL GDNF (Stem cell technologies # 78058), 1μg/mL laminin (Life technologies # 23017-015), 3.3 μg/mL cAMP (Stem cell technologies # 73884), 3.52 μg/mL Ascorbic acid (Stem cell technologies # 72132), 0.5mM L-glutamine and 1% NEAA and then plated on to T25 flasks. iPSC cortical neurons were harvested on DIV 65-72 for RNA sequencing analysis.

### Immunocytochemistry of iPSC-MG

For iPSC-MG DIV 28-33 of microglia differentiation, non-adherent iPSC-MGs were collected and plated onto a 4 or 8 well fibronectin (sigma # F0895; 1:40) coated glass bottom chamber slides (Ibidi # 80427 or #80827) at a seeding density of 250 000 or 125 000 cells per well respectively. One hour after plating, cells were fixed with 4% paraformaldehyde (PFA; Electron Microscopy Sciences # 15714-S) for 20 minutes, washed three times in PBS for 5 minutes and then, blocked with 0.2% Triton X-100 and 5% Normal Goat Serum (Vector # S1000) for 1 h at room temperature. Primary antibodies were prepared in blocking solution and applied overnight at 4 °C. The following primary antibodies were used during our studies: anti-PU.1 (Cell Signaling Technology # 2266S) 1:500; anti-P2RY12 (Sigma # HPA014518) 1:500; anti-CX3CR1 (Biorad/AbD Serotec # AHP1589) 1:500; anti-TREM2 (abcam # AB209814) 1:500; anti-TMEM119 (abcam # ab185333) 1:100; anti-LAMP1 (Developmental Hybridoma Bank # H4A3-s) 1:100; anti-EAA1 (BD Biosciences # 610457) 1:700; anti-C9orf72 (Sigma # HPA023873) 1:100; anti-TDP-43 (Cell signaling # 89789, TDP-43 D9R3L) 1:500; anti-ADAR-2 (Sigma # HPA018277)1:500. Next, cells were washed in PBS three times for 7 min and then, incubated consecutively with respective fluorophores secondary antibodies. Alexa Fluorophores (Invitrogen) were used at a 1:750 and prepared in blocking solution without triton and incubated for 45 minutes at room temperature. Cells were then washed with PBS three times for 7 min each and DAPI was applied. For nuclear markers TDP-43 and ADAR, wheat germ agglutinin 680 (Invitrogen # W32465) was used to label iPSC-MG cell surface. Mounting media ibidi (ibidi # 50001) was used on chambers slides.

### RNA isolation, whole transcriptome library preparation, and sequencing

At DIV 28-30 iPSC-MG from 8 *C9orf72* ALS/FTD patient lines and 4 control lines were pelleted and lysed using QIAshredder (QIAGEN-79654) and RNA was isolated with RNeasy Mini Kit (QIAGEN-74104) following the manufacturer’s instructions. RNA samples were measured for quantity with Quant-iT Ribogreen RNA Assay (Thermo Fisher, Cat. No. R11490) and quality with Agilent High Sensitivity RNA ScreenTape and buffer (Agilent, Cat. No. 5067-5579 & 5067-5580). For each RNA sample, an indexed, Illumina-compatible, double-stranded cDNA whole transcriptome library was synthesized from 1µg of total RNA with Takara Bio’s SMARTer Stranded Total RNA Sample Prep Kit – HI Mammalian (Takara Bio, Cat. No. 634876) and SMARTer RNA Unique Dual Index Kit (Takara Bio, Cat. No. 634418). Library preparation included ribosomal RNA depletion, RNA fragmentation (94 °C for 3 min), cDNA synthesis, and a 12-cycle unique dual indexing enrichment PCR. Each library was measured for size with Agilent’s High Sensitivity D1000 ScreenTape and reagents (Agilent, Cat. No. 5067-5584 & 5067-5603) and concentration with KAPA SYBR FAST Universal qPCR Kit (Kapa Biosystems, Cat. No. KK4824). Libraries were then combined into an equimolar pool which was also measured for size and concentration. The pool was clustered onto a paired-end flowcell (Illumina, Cat. No. 20012861) with a 20% v/v PhiX Control v3 spike-in (Illumina, Cat. No. FC-110-3001) and sequenced on Illumina’s NovaSeq 6000. The first and second reads were each 100 bases. All Aβ (1-40) and LPS stimulated cells at DIV 28-30 iPSC-MG from 4 *C9orf72* ALS/FTD patient lines and 5 control lines were processed as previously described.

### Human tissue RNA sequencing

We accessed human brain tissue RNA sequencing performed by Target ALS and the New York Genome Center (http://www.targetals.org/research/resources-for-scientists/resource-genomic-data-sets/). Sixteen cases of control frontal cortex, 8 *C9orf72* ALS/FTD frontal cortex, 15 control motor cortex, 12 *C9orf72* ALS/FTD frontal cortex, 4 control occipital cortex and 5 *C9orf72* ALS/FTD frontal cortex were evaluated for differential expression of microglia specific genes.

### RNA sequencing analysis

Fastq files were quality and adapter trimmed using cutadapt (version 1.14). Adapter trimmed fastq files were then aligned to the human genome (hg38, gencode v29) using STAR (version 2.6.1d) with default options. RNA count matrices were pulled from aligned BAM files using featureCounts (version 1.6.4). All downstream statistical analysis was done in R (version 3.6.2) using raw counts matrices from featureCounts as input. Low expression genes were filtered such that genes with mean read counts < 10 were removed from the analysis. Differential expression analysis was done using DESeq2 (version 1.26.0) using disease status or treatment regime as the model. Volcano plots were generated from DESeq2 output using EnhancedVolcano. Heatmaps were generated using heatmap from z-scores calculated from DESeq2 normalized gene counts. Tissue data from Target ALS was downloaded from the New York Genome Center as raw fastq files and pushed through an identical analysis pipeline as data generated in our lab.

### Repeat primed PCR to detect the presence of C9 HRE in iPSC and iPSC-MG

We followed previous established protocols (Renton et al., 2011a) to determine the presence the hexanucleotide repeat expansion (G_4_C_2_: >30) in *C9orf72* of iPSC and iPSC-MG (Table S2).

### Real-time quantitative RT-PCR

RNA was isolated using Qiagen RNeasy Micro Kit (Cat #74004) according to manufacturer’s instructions. RNA was reverse transcribed to cDNA with oligo(dT) with the Promega Reverse Transcriptase System (Cat # A3500) and analyzed using SYBR Green Master Mix (Applied Biosystems). *C9orf72* (Forward-5’-CAGTGATGTCGACTCTTTG-3’ and Reverse-5’ AGTAGCTGCTAATAAAGGTGATTTG-3’). Expression was normalized to RPL13A (Forward-5’ CCTGGAGGAGAAGAGGAAAGAGA-3’ and Reverse-5’ TTGAGGACCTCTGTGTATTTGTCAA-3’) or B2M (Forward-5’ TGCTGTCTCCATGTTTGATGTATCT-3’ and Reverse-5’ TCTCTGCTCCCCACCTCTAAGT-3’).

### Western blotting

Microglia cell pellets were homogenized in RIPA lysis and extraction buffer (Thermo Scientific, Cat #89900), supplemented with protease inhibitor cocktail (complete, Roche) and phosphatase inhibitor cocktail (PhosSTOP, Roche). Protein concentration was determined by BCA assay kit (Thermo Fisher, Cat# 23225). Cell lysates were separated on 4-20% protean TGX precast gels (Biorad, Cat # #4561096) and blotted onto nitrocellulose membranes (Biorad, Cat # 1704159). Membranes were blocked for 60 min with Odyssey blocking buffer (PBS, Li-Cor, Cat #927-40000) and incubated overnight at 4°C with anti-C9orf72 (GeneTex Cat #. GTX634482, 1:1000), and anti-actin (Sigma-Aldrich A5441, 1:5000) antibodies. We used bone marrow derived macrophages (BMDM) from a *C9orf72* knockout mouse (O’Rourke et al., 2016) to validate the antibody used for the western blot analysis. After washing membranes were incubated for 60 min with IRDye fluorescent secondary antibodies (Li-Cor). After washing, blots were subsequently analyzed with Li-COR imaging system (Odyssey CLx).

### Repeat hybridization chain reaction (R-HCR) to detect intranuclear repeat RNA foci

We followed previous published methods (Glineburg et al. 2021). All main reagents and probes were purchased from Molecular Instrument, Inc. The negative control probe lacked any binding region with tail ends of the initiator probe that bind to the hairpins. Briefly, on DIV 30 of microglia differentiation, iPSC-MGs were plated on to 8 chambers glass bottom slides (Ibidi #80827) coated with human fibronectin (sigma # F0895; 1:40) at a cell density of 125 000-150 000 cells per chamber. *Fixation step* One hour after plating, iPSC-MG were washed once with 1x PBS and then fixed with 4% PFA for 10 min at room temperature. We further fix cells in 70 % cold ethanol overnight at 4°C. *Initiation/Hybridization step* Next day, 70% cold ethanol was removed and 250 µl per well of preheated hybridization buffer was added to cells and incubated at 45°C for 30 min. The hybridization buffer was then replaced with 125µl of pre-warmed initiation probe and incubated at 45°C overnight (12 – 16 h). 4nM C9 (CCCCGG)_6_ and negative scramble initiator probes solutions were prepared in 45°C pre-warmed hybridization buffer. *Amplification step* Six hours before starting the amplification stage, the R-HCR probe washing buffer was thawed and warmed up at 45°C. Cells were then washed 4x with pre-warmed (45°C) HCR probe wash buffer for 5 mins at 45°C. Then washed 2x for 5 mins with 5x SSC-T (5 X SSC, 0.1 % Tween20) at room temperature. Cells were then left in 5x SSC-T in a humid chamber at room temperature until amplification stage was started. Approximately 1 hour prior to start the amplification stage we thawed the amplification buffer and hairpin probes B1H1 and B1H2 Alexa Fluor-546 to room temperature. Separately, hairpin B1H1 and hairpin B1H2 were prepared by snap cooling to the appropriate volume of the 3 µM stock. 15nM of each hairpin probe was used. Each hairpin was heated up at 95°C for 90 seconds, place on ice and then leave in a dark drawer at room temperature (no ice) for 30 min. We combined the snap-cooled B1H1 hairpins and snap-cooled B1H2 hairpins in the appropriate volume of amplification buffer at room temperature. The pre-amplification solution was removed and 125 µl per well of the hairpin solution was added to each chamber. iPSC-MGs were incubated cells with hairpin probes in the dark at room temperature overnight (12 – 16 hours). Next day, cells were washed x5 for 5 minutes each in 5x SSC-T at room temperature. Then, incubated with DAPI (made in 5x SSC-T) for 10 mins at room temperature. Wash cells 2x for 7 minutes each in 5x SSC-T at room temperature. Mount with Ibidi mounting medium (Ibidi #50001). IPSC-MGs were then visualized and imaged using a Zeiss LSM800 laser scanning confocal microscope. Using Imaris Software from Bitplane, we quantified the percentage of iPSC-MG presenting HRE-associated nuclear foci and the total number of foci per cell.

### Immunoassay analysis of poly-(GP)

Levels of poly-(GP) in cell lysates were measured in a blinded fashion using a Meso Scale Discovery (MSD) immunoassay and a MSD QUICKPLEX SQ120 instrument. A purified mouse monoclonal poly-(GP) antibody was used as both the capture and detection antibody (TALS 828.179, Target ALS Foundation). The capture antibody was biotinylated and used to coat 96-well MSD Small Spot Streptavidin plates, whereas the detection antibody was tagged with an electrochemiluminescent label (MSD GOLD SULFO-TAG). Lysates were diluted to the same protein concentration, and each sample was tested in duplicate. For each well, the intensity of emitted light, which is reflective of poly-(GP) levels and presented as arbitrary units, was acquired upon electrochemical stimulation of the plates.

### IPSC-MG cytokine assay

On DIV 29, iPSC-MGs were plated on to 4 chambers glass slides coated with human fibronectin (sigma # F0895; 1:40) at a cell density of 250 000 cells per chamber. On DIV 30, control and disease iPSC-MGs were treated with LPS (100ng/mL) for 6 h, based on previous studies (Balez et al., 2016, Muffat et al., 2016). The conditioned media was collected and analyzed for selective cytokine/chemokine profile using the U-PLEX Biomarker Group 1 (Human; Mesoscale #K15067L) Multiplex Assay per manufacturer’s protocol.

### LPS and Aβ (1-40) stimulation of iPSC-MG

On DIV 28-30 iPSC-MG from 4 *C9orf72* ALS/FTD patient lines and 5 control lines were plated on to 4 chambers glass slides as described above. For *LPS stimulation* -One hour after plating, iPSC-MGs were treated for 6 h with LPS (100ng/mL). All iPSC-MG were then collected and sent for RNA sequencing analysis. For *Aβ (1-40) stimulation* -iPSC-MGs were treated for 2 h with vehicle (DMSO) or 1µM Aβ (1-40) TAMRA (human Aβ, AnaSpec # AS-60488; (Paolicelli et al., 2017)) prepared in microglia basal media with growth factors. All iPSC-MGs were collected at 2 hours and sent for RNA sequencing analysis.

### Phagocytosis of Aβ (1-40) TAMRA protein by iPSC-MG

On DIV 30 of microglia differentiation, iPSC-MGs were plated on to 4 chambers glass slides coated with human fibronectin (sigma # F0895; 1:40) at a cell density of 250 000 cells per chamber. One hour after plating, iPSC-MGs were treated for 5 minutes with vehicle (DMSO) or 1µM of fluorescently labeled Aβ (1-40) TAMRA (human Aβ, AnaSpec # AS-60488; (Paolicelli et al., 2017)) prepared in microglia basal media with growth factors. All iPSC-MGs were washed at 5min with microglia basal media supplemented with growth factors. IPSC-MGs were then fixed after 5 minutes, 30 minutes, and 1 h and immunostained for TREM2 as described above. Only 5 minutes and 1 h time points were quantified. Cells were imaged using a Zeiss LSM800 confocal microscope. Using Imaris Software from Bitplane we determine the cell volume and percentage of the microglia surface area covered by Aβ (1-40) TAMRA at different time points.

### Engulfment of human brain synaptoneurosomes by iPSC-MG

Fluorescently labeled pHrodo (a pH sensitive dye that fluoresces only in acidic compartments) human brain synaptoneurosomes (hSN-rodo) were generated as described previously (Hesse et al., 2019, Tzioras et al., 2019). On DIV 30 of the microglia differentiation, iPSC-MGs were plated on to 8 chamber glass slides that were coated with human fibronectin (sigma # F0895; 1:40) at a cell density of 100 000 cells per chamber. One hour after plating, we labeled live microglia with the nuclear marker Hoechst 33342 (Thermo fisher # H3570). Then, control and *C9orf72* iPSC-MGs were treated with 1:100 dilution of 4mg/mL hSN-rodo in the presence or absence of 10 µM cytochalasin-D (inhibitor of actin polymerization). Confocal live cell imaging of iPSC-MGs was done using a 20X objective of a Zeiss 800 confocal microscope. IPSC-MGs were imaged every 10 minutes for up to 6 h for initial phagocytosis studies to determine the percentage of cells that exhibit phagocytic activity. For all other set of experiments, iPSC-MGs were imaged every 10 minutes for up to 2 h. All lines had three technical replicates for the hSN-rodo treatment. Six images were taken per well per line for a total of 18 images every 10 minutes. All images were analyzed using Imaris Software from Bitplane. To determine the percentage of iPSC-MGs engulfing hSN-rodo, the spots module was used to count the total number of iPSC-MGs based on Hoechst marker and the total number of phagocytic iPSC-MGs per image. Additionally, we quantified the fluorescence mean intensity of the cargo hSN-rodo per phagocytic cell at the 2 h time point, where more than 60% of the iPSC-MGs were engulfing hSN-rodo. Cytochalasin-D was used as a negative control.

### Confocal microscopy and bright-field imaging

All immunostained iPSC-MGs were visualized and imaged using a Zeiss LSM800 laser scanning confocal microscope. Per staining, all images were taken with same settings for parallel cultures. For all iPSC-MG immunostainings and Aβ (1-40) TAMRA phagocytic activity assay, a plan Apochromat 63x oil immersion objective was used; Z-stacks were generated with 1024 x 1024 image size, 0.5x XY scan zoom and 1µm scaling. For some immunostainings, differential interference contrast (DIC) was used to highlight iPSC-MG surface area. For live cell imaging of iPSC-MG engulfing synaptoneurosomes, confocal microscopy with differential interference contrast was used. Tiled images were captured using a 20X objective with a 1.0x XY scan zoom and 0.624 µm x 0.624µm scaling. Bright-field images of the iPSC-MG cultures were taken using a Zeiss AxioVert.A1 microscope and a resolve HD Ludesco camera.

### Imaging analysis using Imaris Software from Bitplane

All images were processed and analyzed using Imaris Software 9.5.1 and 9.6 from Bitplane. In order to obtain volume, area, mean intensity and sum intensity per cell for a large number of samples, we assigned a randomized color identification per cell followed by the use of ImarisVantage module to extract multiple numerical values from the created 3 dimensional structures. For *iPSC-MG marker characterization*, the spots module was used to count the total number of cells positive for a specific microglia marker per image while using the DAPI channel as a reference. To calculate the *TDP-43 and ADAR-2 nucleocytoplasmic ratio (N/C ratio),* we used the surface module to generate iPSC-MG microglia 3 dimensional cellular (based on membrane staining with wheat germ agglutinin) and nuclear surfaces (based on DAPI). The sum intensity of TDP-43 or ADAR-2 was acquired for the nucleus and cytoplasm as well as the volume for each cellular compartment. The following formula was used: (Cell Sum Intensity – Nucleus Sum Intensity) = Cytoplasm Sum Intensity; (Cell Volume – Nucleus Volume) = Cytoplasm Volume. *N/C ratio* = (Nucleus Sum Intensity/ Nucleus volume) / (Cytoplasm Sum Intensity/ Cytoplasm volume). Data from each cell was acquired by assigning randomized color identification followed by the use of ImarisVantage. For the *Aβ (1-40) TAMRA phagocytic activity*, we used *Aβ (1-40) TAMRA* fluorescence signal and chose an ideal threshold to create the *Aβ* surfaces inside the IPSC-MGs. TREM2 staining was used to generate the cell surface structures. An algorithm was also generated and used in all iPSC-MG parallel cultures. Randomized color identification and ImarisVantage was also used to extract all data like previous data sets. For *hSN-rodo live cell imaging over time,* the spots module was used to count the total number of cells over time. Phagocytic cells were manually identified based on hSN-rodo signal inside iPSC-MG. We set a threshold between 10,000-12,000 units above grey scale as the positive signal indicative of engulfment and lysosomal internalization. At the 2h time point, the cell surface area of phagocytic cells was manually outlined using Imaris Software manual creation tool and the hSN-rodo mean intensity value was obtained per cell. For *iPSC-MGs EEA1 and Lamp1 analysis,* based on the staining pattern we manually choose an optimal threshold for each protein marker to generate the 3D surfaces. We then, stored the surfaces algorithms and used it in all iPSC-MG parallel cultures. Randomized color identification and ImarisVantage were then used to extract the cell volume and mean intensity of both markers per cell.

### Statistical analysis

Statistical Analysis for RNA sequencing was done in R (version 3.6.2) and detailed above. All other statistical analyses were performed using Graphpad Prism 7 and 8. For comparison of two groups we used two-tailed Student’s t-test. Two tailed Mann Whitney test was performed for the poly-(GP) ELISA assay. All statistical significance was ranked as the following: * *p* ≤ 0.05; ** *p* ≤ 0.01; *** *p* ≤ 0.001; **** *p* ≤ 0.0001 and *p* > 0.05 not significant. All other statistical details and exact p-values are reported in each Figure legend.

### Data availability (accession numbers)

All RNA sequencing data, and analysis code used in this publication are available via accession number syn22335553.

## Acknowledgments

We are grateful for all human tissue samples donated to Target ALS to be made available for our research. We are also thankful for all patients that donated cells for the generation of iPSCs to be used in our studies. We further acknowledge the receipt of iPSC lines through the ‘Cedars-Sinai Medical Center’s David and Janet Polak Foundation Stem Cell Core Laboratory’. We thank both the Target ALS Consortium and the New York Genome Center for access to their RNA sequencing database. In particular, we would like to thank Drs. Lyle Ostrow and Hemali Phatnani. We would also like to acknowledge Dr. Ramita D. Karra and Dr. Bryan J. Traynor for their technical expertise. We would like to give special thanks to members of the Sattler laboratory for their suggestions and feedback on the manuscript. We would also like to give thanks to Dr. Ileana Soto-Reyes and Dr. Nadine Bakkar for scientific advice and feedback on the manuscript. This research was supported by the National Institute on Aging of the National Institutes of Health on the Award Number P30AG019610 (RS) and R01NS120331 (RS and KKJ). The content is solely the responsibility of the authors and does not necessarily represent the official views of the National Institute of Health. This work was further supported in part by a grant to Cedars-Sinai Medical Center from the Howard Hughes Medical Institute through the James H. Gilliam Fellowships for Advanced Study program. In addition, support was provided through funds by The Robert Packard Center for ALS Research (RS), Barrow Neurological Foundation (RS; RB), The David E. Reese Family Foundation and the George Flaccus Laboratory at TGen for ALS Research (KKJ), Barrow Neurological Foundation postdoctoral fellowship (IL), The Fein Family Foundation (RS; RB), U.S. Department of Veterans Affairs I01BX003625 (RS), NIH/NINDS RO1NS101986 (FBG), NIH/NINDS, RO1NS097545 (RHB), NIH/NINDS T32 NS082174 (AM).

## Author contributions

IL: Designed experiments, differentiated iPSC-MG, performed and supervised experiments; performed quantitative confocal microscope image analysis; acquired and analyzed data and wrote the manuscript. EA: Performed RNA sequencing analysis for iPSC-MG and human postmortem tissue. JL: Differentiated and maintained iPSC-MG. LMG, BER, DB: Acquired data and performed quantitative microscope image analysis. DL, LB, JS, TAM, TFG: Performed and analyzed experiments. MT, AL, JR: Performed experiments. SM, DB: Performed confocal imaging. RP, EH: RNA sequencing analysis. MS, CJD, MM: Performed quantitative image analysis. CB: Maintained and expanded iPSCs. FBG, SA, JI, MH: Provided *C9orf72* iPSC lines. MBJ, AM: Taught first-hand iPSC-MG protocol. TSJ: Provided hSN-rodo. RB, RHB: Provided suggestions and evaluation of our work. KKJ: Designed experiments, provided critical advice and evaluation of our work. RS: Designed experiments, oversaw data analysis and interpretation and participated in manuscript writing and editing.

## Figure Legends for Supplementary Figures

**Supplementary Figure 1.**
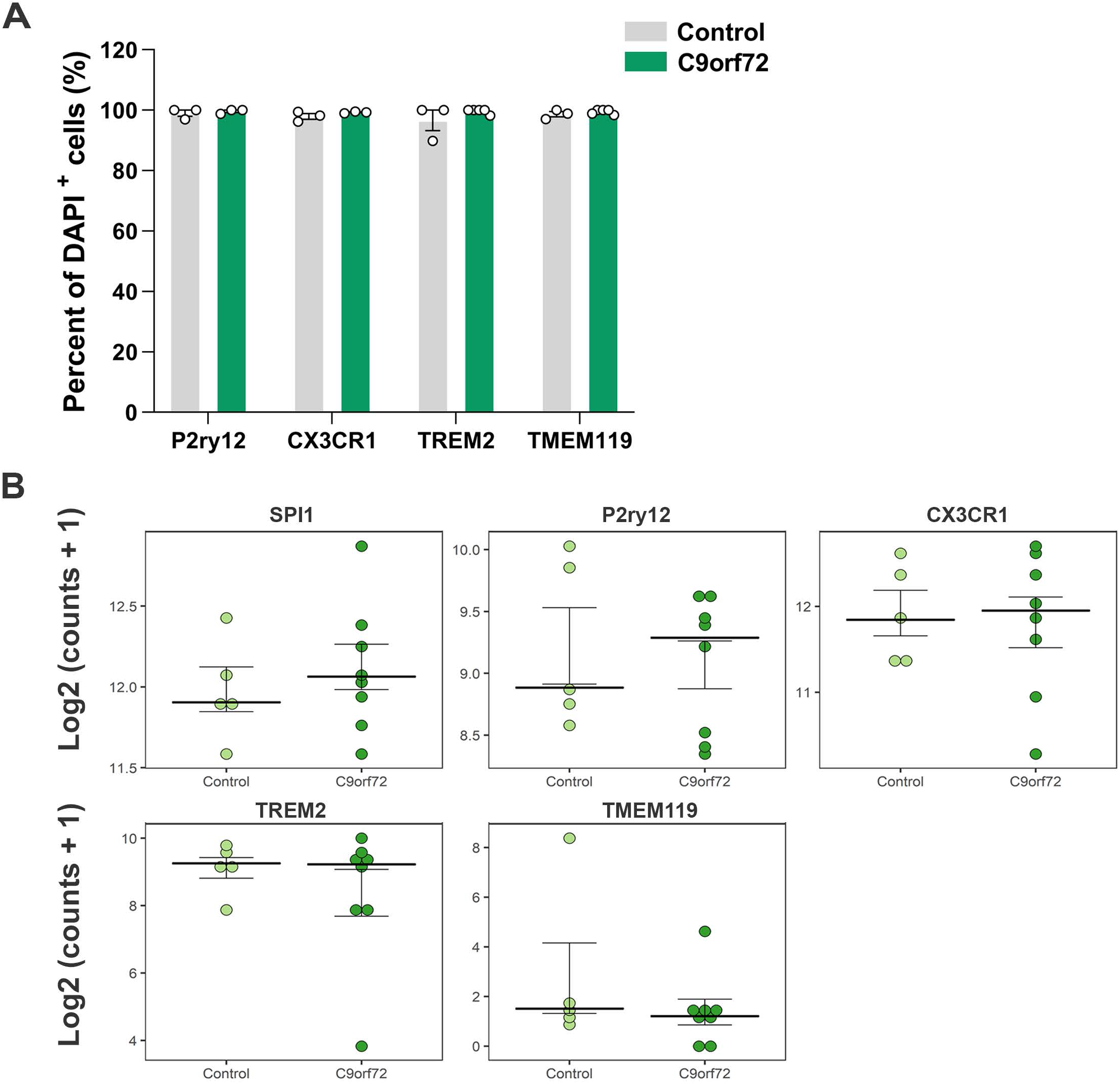
Expression of microglia specific genes and microglia protein markers in healthy control and *C9orf72* ALS/FTD iPSC-derived microglia. (A) Percentage of DAPI-positive cells from mature healthy control (n=3 lines, 1-2 differentiations per line) and *C9orf72* ALS/FTD (n=3-5 lines, 1-2 differentiations per line) iPSC-MG positive for P2ry12, Cx3cr1, TREM2 and TMEM119 protein. (B) Dot plots showing the level of expression as Log2 (counts +1) of microglia specific genes *spI1* (encodes for myeloid transcription factor PU.1), *P2RY12*, *CX3CR1*, *TREM2* and *TMEM119* in healthy control and *C9orf72* ALS/FTD (Control, n=4 lines, 1-2 differentiations per line; *C9orf72*, n=7 lines, 1-2 differentiations per line).

**Supplementary Figure 2.**
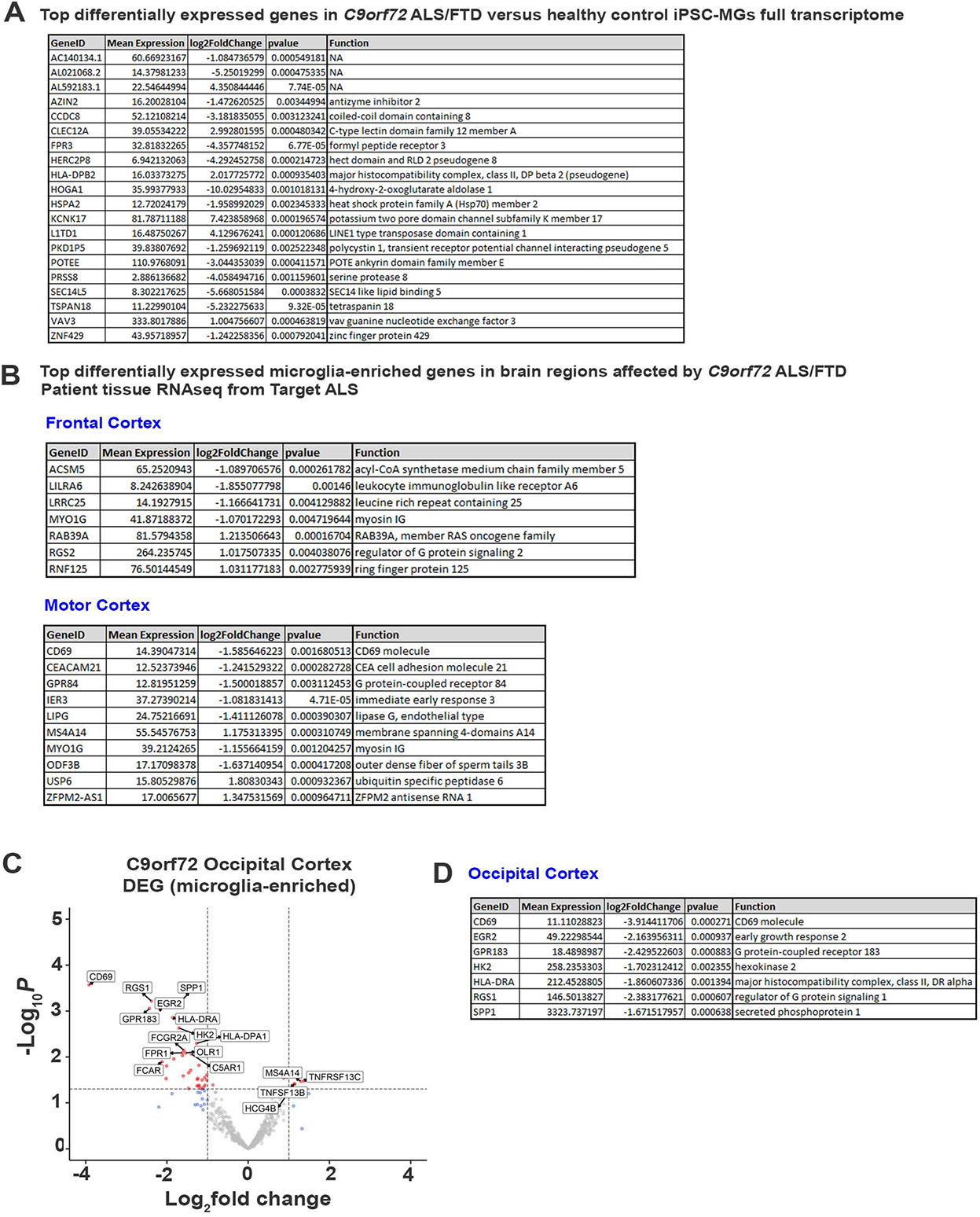
Differentially expressed transcripts in *C9orf72* ALS/FTD versus healthy control iPSC MG full transcriptome and dysregulated microglial-enriched transcripts in *C9orf72* ALS/FTD frontal, motor and occipital cortex. (A) Top differentially expressed transcripts in iPSC MG full transcriptome (unadjusted p value <0.005; log_2_ fold change (FC) ± 1). (B) Top microglial-enriched dysregulated transcripts in the frontal and motor cortex of *C9orf72* ALS/FTD patient tissue from bulk RNA sequencing. (C) Volcano plot of differentially expressed microglial-enriched genes (total of 881 from (Gosselin et al., 2017)) in *C9orf72* ALS/FTD occipital cortex (control, n=4 lines; *C9orf72*, n=5 lines) (unadjusted p value <0.005; log_2_ fold change (FC) ± 1). (D) Dysregulated transcripts in the occipital cortex of *C9orf72* ALS/FTD patient tissue from bulk RNA sequencing. All human brain RNA sequencing data was taken from Target ALS/NYGC.

**Supplementary Figure 3.**
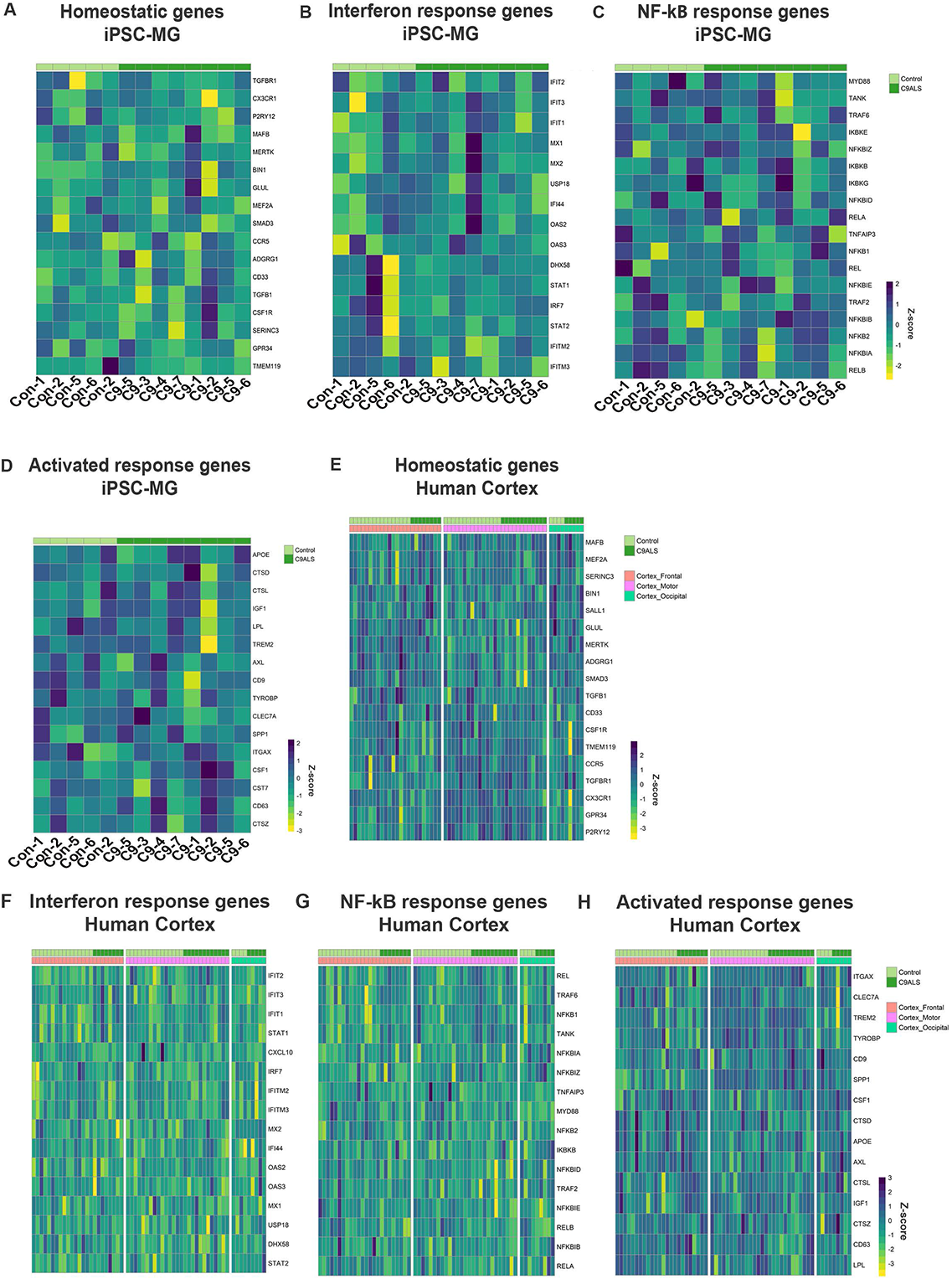
Heatmap of the homeostatic, interferon, NF-kB and activated response genes in iPSC-MG and human cortex. Expression of (A, E) homeostatic, (B-F) interferon, (C-G) NF-kB and (D-H) activated response genes in iPSC-MGs (control, n=4 cell lines with 1-2 differentiations each and *C9orf72* ALS/FTD, n=7 cell lines with 1-2 differentiations each) and human cortex (frontal cortex control n=16 and *C9orf72* ALS/FTD n=8; motor cortex control n=15 and *C9orf72* ALS/FTD n=12 and occipital cortex control n=4 and *C9orf72* ALS/FTD n=5).

**Supplementary Figure 4.**
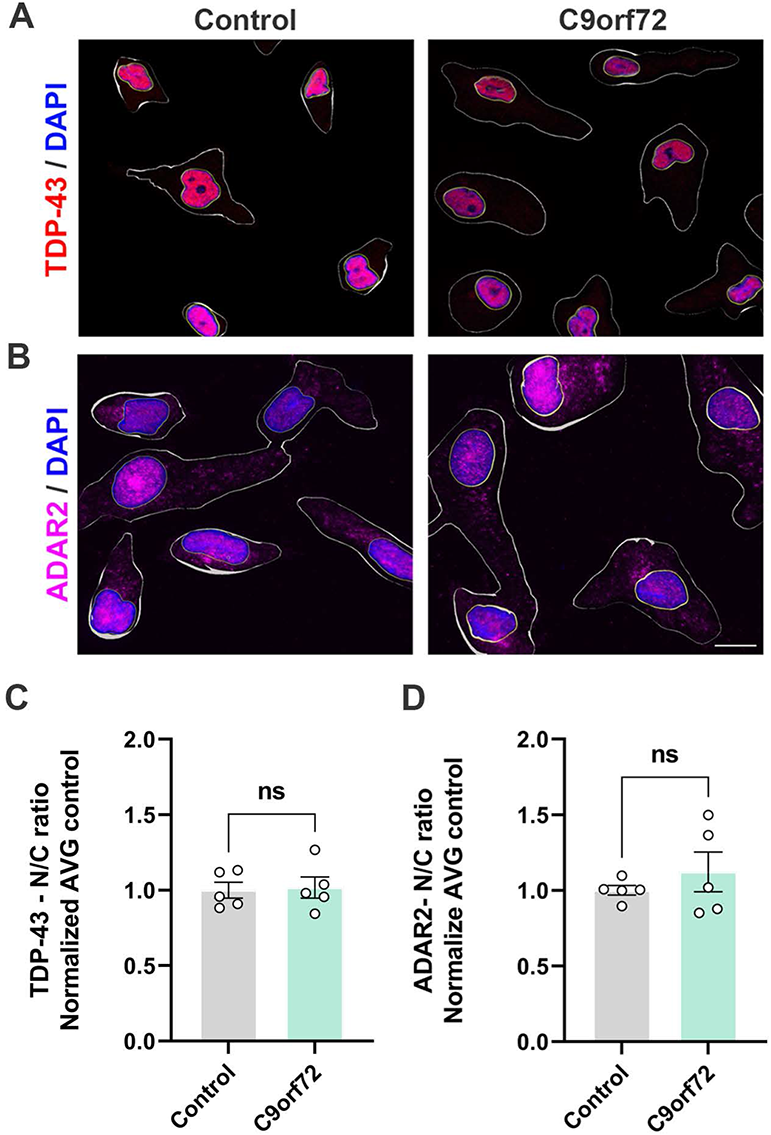
*C9orf72* ALS/FTD iPSC-MG monocultures do not exhibit TDP-43 pathology or cytoplasmic mislocalization of ADAR2. (A) TDP-43 nuclear staining in control and *C9orf72* ALS/FTD iPSC-MG. IPSC-MG cell surface and nuclear surface is outlined (white). Scale bar, 15µm. (B) Control and *C9orf72* ALS/FTD iPSC-MG immunostained for anti-ADAR2. IPSC-MG nuclear and cell surface is outlined (white). Scale bar, 10µm. (C) Quantification of TDP-43 nucleocytoplasmic ratio using Imaris Software. No evidence of cytoplasmic accumulations in *C9orf72* ALS/FTD iPSC-MG monocultures (control, n=5 lines; *C9orf72*, n=5 lines, n=60-84 cells per line; p= 0.85, Student’s t-test). (D) Quantification of ADAR2 nucleocytoplasmic ratio. No evidence of nuclear ADAR2 mislocalization to the cytoplasm was observed in *C9orf72* ALS/FTD iPSC-MG monocultures (control, n=5 lines; *C9orf72*, n=5 lines, n=100-115 cells per line; p=0.39, Student’s t-test).

**Supplementary Figure 5.**
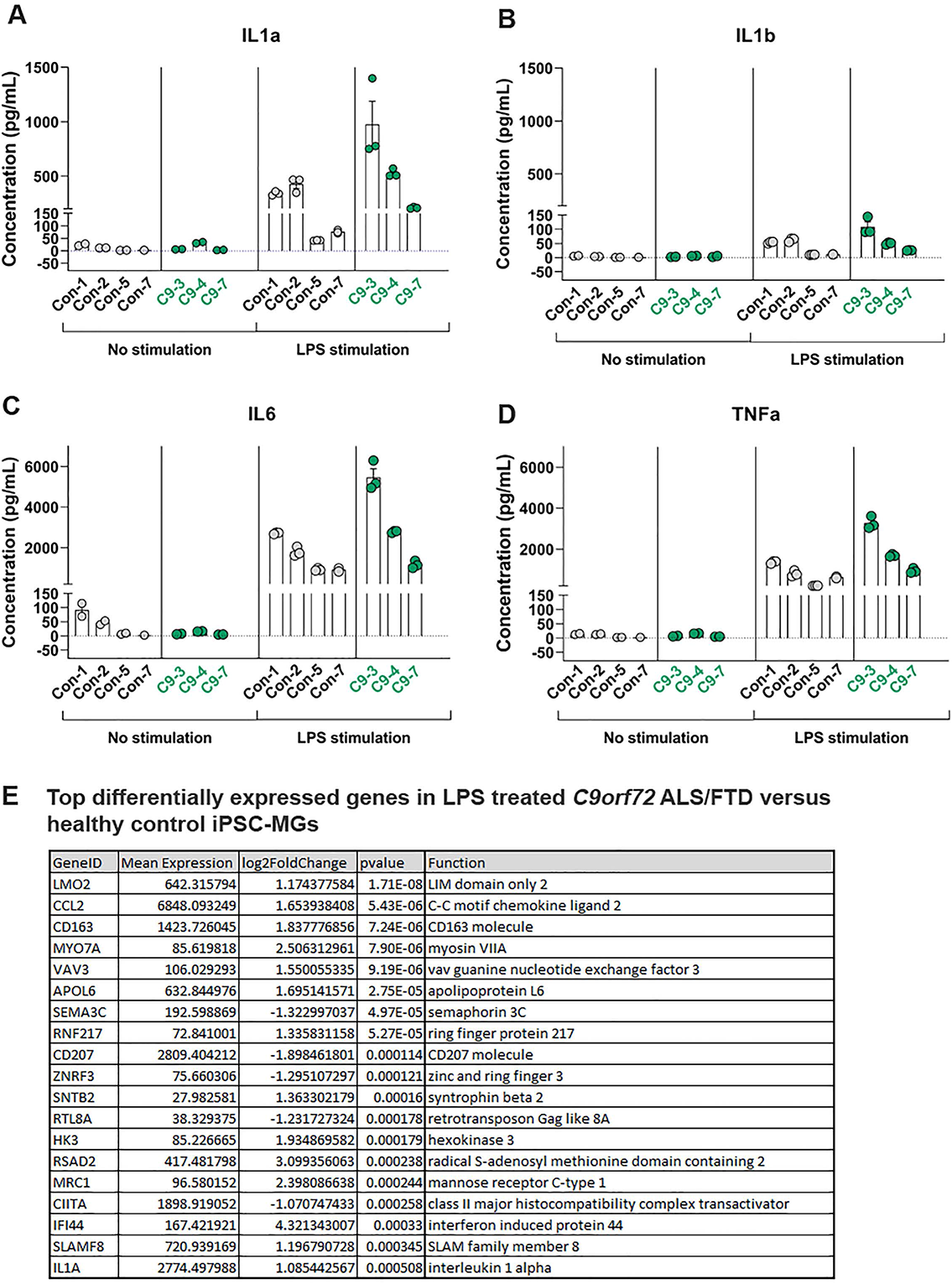
*C9orf72* ALS/FTD iPSC-MG response to LPS stimulation. Control IPSC-MGs (n=4) and *C9orf72* ALS/FTD iPSC-MG (n=3) were treated with LPS (100ng/ml) for 6hr. Conditioned media samples were collected and measured by a U-plex Biomarker Group 1 (Human, Mesoscale) Multiplex Assay ELISA to obtain a cytokine/chemokine profile of iPSC-MG. Here we present the concentration (pg/mL) of (A) IL1-α, (B) IL-1β, (C) IL6, and (D) TNF-α per replicate within a sample. (E) Top differentially expressed genes in LPS treated *C9orf72* ALS/FTD versus healthy control iPSC-MGs.

**Supplementary Figure 6.**
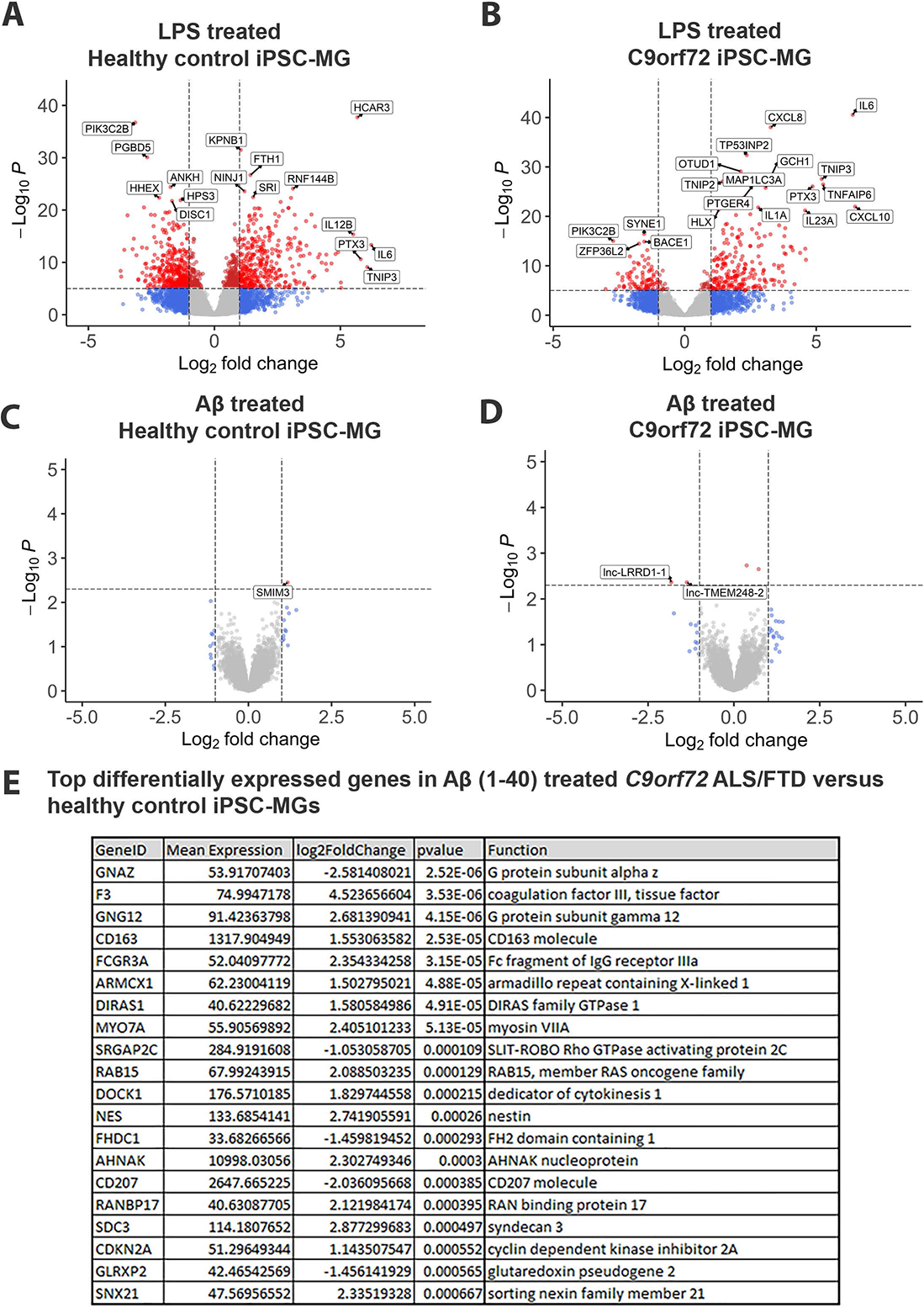
Top differentially expressed genes in LPS or Aβ (1-40) treated healthy control and *C9orf72* ALS/FTD iPSC-MGs. On DIV 28-30, control (n = 5) and *C9orf72* ALS/FTD (n= 4) iPSC-MG were treated for 6 h with LPS (100ng/mL) or for 2 h with 1µM Aβ (1-40) TAMRA. All iPSC-MGs were collected after stimulation and sent for RNA sequencing analysis. Significant dysregulated genes (log_2_ fold change (FC) ± 1, p value <0.05) were observed in both LPS treated healthy controls (A) and LPS treated *C9orf72* ALS/FTD iPSC-MGs (B). Minimal dysregulated genes were observed for all iPSC-MG treated with Aβ (1-40) TAMRA (C-D).

**Supplementary Figure 7.**
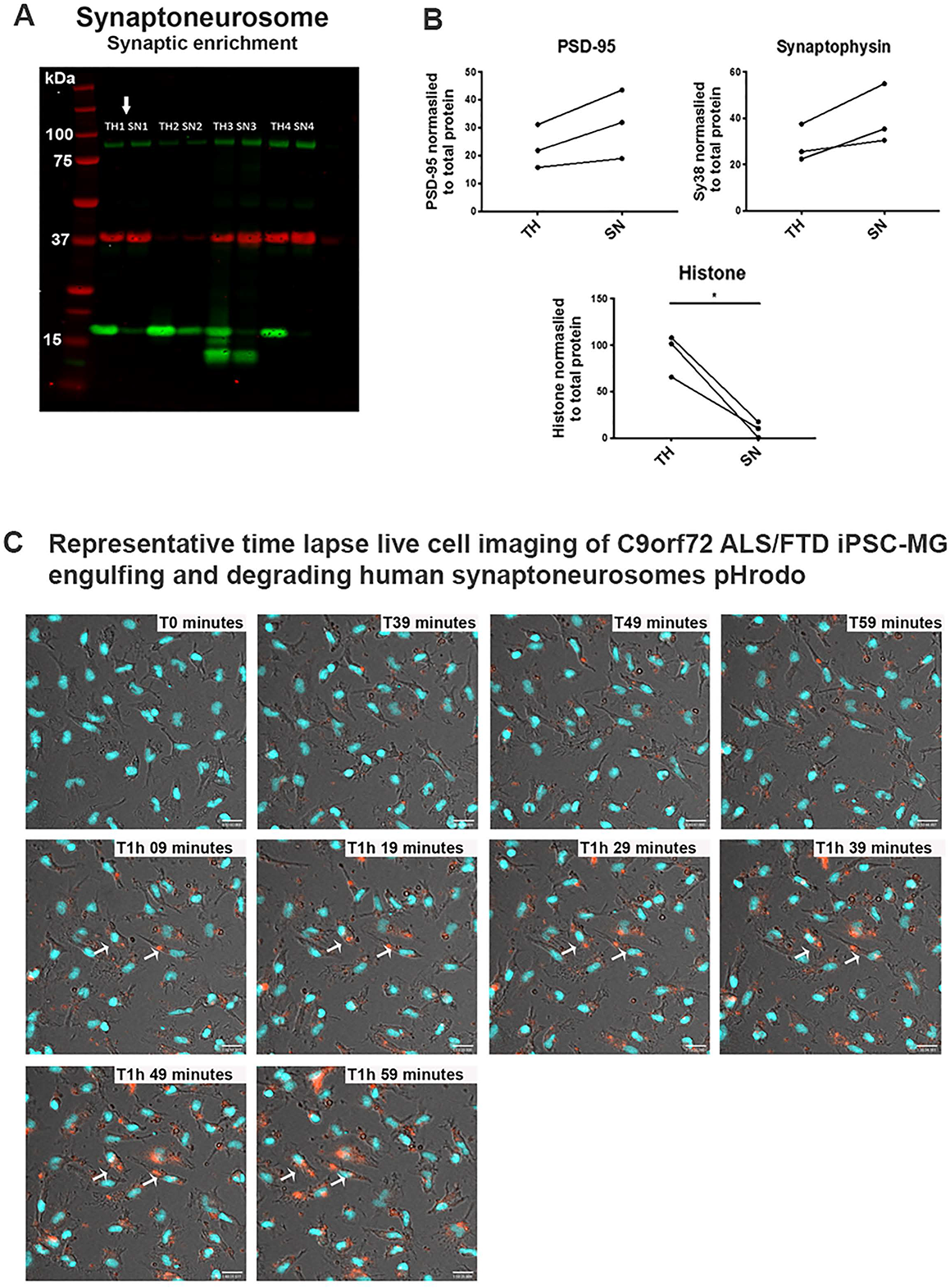
*C9orf72* ALS/FTD iPSC-MG engulf and degrade human pHrodo synaptoneurosomes containing PSD-95 and synaptophysin 38. (A) Human brain pHrodo synaptoneurosomes (hSN-rodo) contain PSD95 and Synaptophysin 38 proteins from pre- and post-synaptic compartments. Western blot analysis for total homogenate (TH) and synaptoneurosome preparations (SN). Four different SN preparations (TH1-TH4 and SN1-SN4) from four different control brains frozen samples. Preparations 1, 3 and 4 had successful enrichment of post-synaptic density 95 (PSD95 – 95kDa) and synaptophysin (Synaptophysin 38kDa) compared to total homogenate. In addition, low levels of nuclear marker histone 3 were observed in the SN preparations 1, 3, 4. SN1 (white arrow; TH1 and SN1) was the preparation used for the present study. (B) PSD95, synaptophysin 38 and Histone 3 quantification for TH and SN preparations 1, 3 and 4. A significant decrease in Histone 3 levels is observed in all SN preparations, p-value =0.02, Student’s t-test. (C) Selected representative images of the time lapse live cell imaging of *C9orf72* ALS/FTD iPSC-MG engulfing and degrading human synaptoneurosomes pHrodo (hSN-rodo). IPSC-MGs were labeled with the live nuclear marker Hoechst (blue) to identify individual cells followed by treatment with hSN-rodo (red). Fluorescent live cell imaging with differential interference contrast microscopy was performed over a 2 h time frame with images taken every 10 min. Here, we present selected images during the T0-T2h time point where more than 60% of iPSC-MGs engulf synapses. HSN-rodo are engulfed rapidly (white arrows highlight several phagocytic iPSC-MGs) and an increase in hSN-rodo intensity is observed in individual cells indicating the uptake of hSN-rodo into acidic intracellular compartments.

**Table S1.** Demographics for *C9orf72* ALS/FTD patient and control iPSCs.

| iPSC line | Gene Mutation | C9 Expansion Size | Gender | Age at disease onset (yrs) | Age at biopsy/blood draw (yrs) | Age at death (yrs) | Disease duration (yrs) | C9 Expansion Size | Comments |
| --- | --- | --- | --- | --- | --- | --- | --- | --- | --- |
| C9orf72 -1 | C9orf72 | >30 repeats | Female | 60 | 63 | N/A | N/A | N/A |  |
| C9orf72 -2 | C9orf72 | >30 repeats | Female | 50 | 51 | 52 | 2 | N/A |  |
| C9orf72 -3 | C9orf72 | >30 repeats | * | * | * | N/A | N/A | ~590 | Almeida et al, Acta Neuropathol, 2013 |
| C9orf72 -4 | C9orf72 | >30 repeats | * | * | * | N/A | N/A | ~200, ~500 | Lopez-Gonzalez et al, Neuron, 2016 |
| C9orf72 -5 | C9orf72 | >30 repeats | * | * | * | N/A | N/A | ~1200 | Almeida et al, Acta Neuropathol, 2013 |
| C9orf72 -6 | C9orf72 | >30 repeats | Male | 52 | 58 | N/A | N/A | N/A |  |
| C9orf72 -7 | C9orf72 | >30 repeats | Male | 48 | 51 | 59 | 11 | N/A |  |
| C9orf72 -8 | C9orf72 | >30 repeats | Male | 57 | 58 | 59 | 1.7 | N/A |  |
| Control 1 | Control | N/A | Female | N/A | N/A | N/A | N/A | N/A |  |
| Control 2 | Control | N/A | Male | N/A | N/A | N/A | N/A | N/A |  |
| Control 3 | Control | N/A | Male | N/A | N/A | N/A | N/A | N/A |  |
| • Control 4 | Control Isogenic | N/A | * | * | * | N/A | N/A | N/A |  |
| Control 5 | Control | N/A | Female | N/A | N/A | N/A | N/A | N/A |  |
| Control 6 | Control | N/A | Male | N/A | N/A | N/A | N/A | N/A |  |
| Control 7 | Control | N/A | Male | N/A | N/A | N/A | N/A | N/A |  |

\* - this information is not provided to protect the ID of the donors
• C9orf72-4 isogenic pair

**Table S2.** Presence of *C9orf72* HRE in iPSC and iPSC-MG cells.

|  |  |  | Repeat primed PCR |
| --- | --- | --- | --- |
| Cell Line | Gene mutation | Cell Type | C9 HRE (>30) |
| C9orf72-1 | C9orf72 | iPSC | Yes |
|  |  | iPSC-MG | Yes |
| C9orf72-2 | C9orf72 | iPSC | Yes |
|  |  | iPSC-MG | Yes |
| C9orf72-3 | C9orf72 | iPSC | Yes |
|  |  | iPSC-MG | Yes |
| C9orf72-4 | C9orf72 | iPSC | Yes |
|  |  | iPSC-MG | Yes |
| C9orf72-5 | C9orf72 | iPSC | not determine |
|  |  | iPSC-MG | not determine |
| C9orf72-6 | C9orf72 | iPSC | Yes |
|  |  | iPSC-MG | Yes |
| C9orf72-7 | C9orf72 | iPSC | Yes |
|  |  | iPSC-MG | not determine |
| C9orf72-8 | C9orf72 | iPSC | not determine |
|  |  | iPSC-MG | not determine |
| Control 1 | Control | iPSC | No |
|  |  | iPSC-MG | No |
| Control 2 | Control | iPSC | not determine |
|  |  | iPSC-MG | not determine |
| Control 3 | Control | iPSC | No |
|  |  | iPSC-MG | No |
| Control 4 | Control Isogenic | iPSC | No |
|  |  | iPSC-MG | No |
| Control 5 | Control | iPSC | not determine |
|  |  | iPSC-MG | not determine |
| Control 6 | Control | iPSC | No |
|  |  | iPSC-MG | not determine |
| Control 7 | Control | iPSC | not determine |
|  |  | iPSC-MG | not determine |

**Table S3.** List of iPSC lines used for specific experiments.

| Cell Line | Gene mutation | Microglia markers | RNAseq | C9 QPCR | C9 WBlot | GP response | TDP43 | ADAR2 | MSD cytokine analysis | Phagocytic activity Aβ (1-40) | hSN Assay hSN-rodo |
| --- | --- | --- | --- | --- | --- | --- | --- | --- | --- | --- | --- |
| C9orf72-1 | C9orf72 | ✓ | ✓ | ✓ | ✓ | ✓ | ✓ | ✓ |  | ✓ | ✓ |
| C9orf72-2 | C9orf72 | ✓ | ✓ | ✓ | ✓ | ✓ |  |  |  | ✓ |  |
| C9orf72-3 | C9orf72 | ✓ | ✓ | ✓ | ✓ | ✓ | ✓ | ✓ | ✓ | ✓ | ✓ |
| C9orf72-4 | C9orf72 | ✓ | ✓ | ✓ | ✓ | ✓ | ✓ | ✓ | ✓ | ✓ | ✓ |
| C9orf72-5 | C9orf72 | ✓ | ✓ | ✓ | ✓ | ✓ |  |  |  | ✓ | ✓ |
| C9orf72-6 | C9orf72 | ✓ | ✓ | ✓ |  | ✓ | ✓ | ✓ |  | ✓ | ✓ |
| C9orf72-7 | C9orf72 |  | ✓ |  |  |  | ✓ | ✓ | ✓ | ✓ | ✓ |
| C9orf72-8 | C9orf72 |  | ✓ |  |  | ✓ |  |  |  |  |  |
| Control 1 | Control | ✓ | ✓ | ✓ | ✓ | ✓ | ✓ | ✓ | ✓ | ✓ | ✓ |
| Control 2 | Control | ✓ | ✓ | ✓ | ✓ |  | ✓ | ✓ | ✓ | ✓ | ✓ |
| Control 3 | Control | ✓ |  | ✓ | ✓ | ✓ |  |  |  | ✓ | ✓ |
| Control 4 | Control Isogenic |  |  | ✓ | ✓ | ✓ |  |  |  |  |  |
| Control 5 | Control | ✓ | ✓ |  |  |  | ✓ | ✓ | ✓ | ✓ | ✓ |
| Control 6 | Control |  | ✓ |  |  |  |  |  |  | ✓ | ✓ |
| Control 7 | Control |  |  |  |  |  | ✓ | ✓ | ✓ |  | ✓ |

**Table S4.** Postmortem tissue sample IDs used for bulk RNA-seq analysis from Target ALS collection.

| ID | Tissue of Origin | Classification |
| --- | --- | --- |
| CGND-HRA-00251 | Cortex_Motor | Control |
| CGND-HRA-00218 | Cortex_Motor | Control |
| CGND-HRA-00590 | Cortex_Motor | Control |
| CGND-HRA-00224 | Cortex_Motor | Control |
| CGND-HRA-00654 | Cortex_Motor | Control |
| CGND-HRA-00569 | Cortex_Motor | Control |
| CGND-HRA-00595 | Cortex_Motor | Control |
| CGND-HRA-00091 | Cortex_Motor | Control |
| CGND-HRA-00568 | Cortex_Motor | Control |
| CGND-HRA-00063 | Cortex_Motor | Control |
| CGND-HRA-01186 | Cortex_Motor | Control |
| CGND-HRA-00594 | Cortex_Motor | Control |
| CGND-HRA-00655 | Cortex_Motor | Control |
| CGND-HRA-00596 | Cortex_Motor | Control |
| CGND-HRA-01187 | Cortex_Motor | Control |
|  |  | 15 |
| CGND-HRA-00134 | Cortex_Motor | C9ALS |
| CGND-HRA-00631 | Cortex_Motor | C9ALS |
| CGND-HRA-00358 | Cortex_Motor | C9ALS |
| CGND-HRA-00321 | Cortex_Motor | C9ALS |
| CGND-HRA-00287 | Cortex_Motor | C9ALS |
| CGND-HRA-00286 | Cortex_Motor | C9ALS |
| CGND-HRA-00359 | Cortex_Motor | C9ALS |
| CGND-HRA-00254 | Cortex_Motor | C9ALS |
| CGND-HRA-00320 | Cortex_Motor | C9ALS |
| CGND-HRA-00288 | Cortex_Motor | C9ALS |
| CGND-HRA-00088 | Cortex_Motor | C9ALS |
| CGND-HRA-00060 | Cortex_Motor | C9ALS |
|  |  | 12 |

Frontal Cortex
| ID | Tissue of Origin | Classification |
| --- | --- | --- |
| CGND-HRA-01179 | Cortex_Frontal | Control |
| CGND-HRA-00212 | Cortex_Frontal | Control |
| CGND-HRA-01379 | Cortex_Frontal | Control |
| CGND-HRA-00751 | Cortex_Frontal | Control |
| CGND-HRA-00021 | Cortex_Frontal | Control |
| CGND-HRA-00476 | Cortex_Frontal | Control |
| CGND-HRA-01236 | Cortex_Frontal | Control |
| CGND-HRA-00242 | Cortex_Frontal | Control |
| CGND-HRA-01225 | Cortex_Frontal | Control |
| CGND-HRA-00529 | Cortex_Frontal | Control |
| CGND-HRA-01234 | Cortex_Frontal | Control |
| CGND-HRA-01228 | Cortex_Frontal | Control |
| CGND-HRA-00466 | Cortex_Frontal | Control |
| CGND-HRA-01185 | Cortex_Frontal | Control |
| CGND-HRA-00788 | Cortex_Frontal | Control |
| CGND-HRA-00871 | Cortex_Frontal | Control |
|  |  | 16 |
| CGND-HRA-00018 | Cortex_Frontal | C9ALS |
| CGND-HRA-00245 | Cortex_Frontal | C9ALS |
| CGND-HRA-00362 | Cortex_Frontal | C9ALS |
| CGND-HRA-00550 | Cortex_Frontal | C9ALS |
| CGND-HRA-00323 | Cortex_Frontal | C9ALS |
| CGND-HRA-00244 | Cortex_Frontal | C9ALS |
| CGND-HRA-00248 | Cortex_Frontal | C9ALS |
| CGND-HRA-00460 | Cortex_Frontal | C9ALS |
|  |  | 8 |

Occipital Cortex
| ID | Tissue of Origin | Classification |
| --- | --- | --- |
| CGND-HRA-01204 | Cortex_Occipital | Control |
| CGND-HRA-01343 | Cortex_Occipital | Control |
| CGND-HRA-01235 | Cortex_Occipital | Control |
| CGND-HRA-00567 | Cortex_Occipital | Control |
|  |  | 4 |
| CGND-HRA-00551 | Cortex_Occipital | C9ALS |
| CGND-HRA-01337 | Cortex_Occipital | C9ALS |
| CGND-HRA-00363 | Cortex_Occipital | C9ALS |
| CGND-HRA-00540 | Cortex_Occipital | C9ALS |
| CGND-HRA-00324 | Cortex_Occipital | C9ALS |
|  |  | 5 |

**Table S5.**
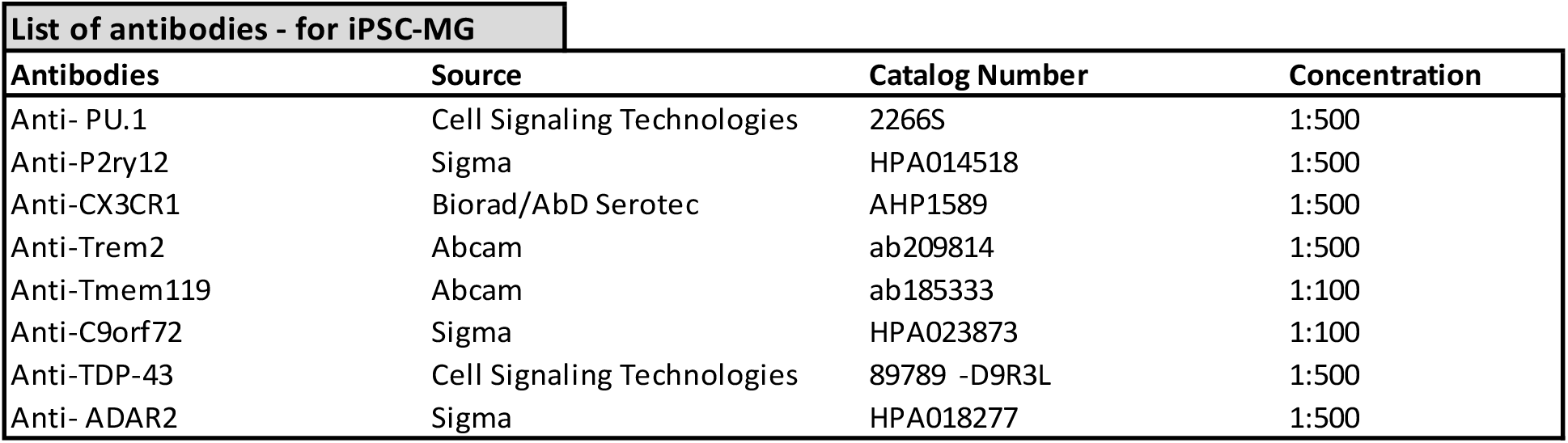
List of antibodies used to stain iPSC-MG cells.

**Movie S1. Time lapse live cell imaging of *C9orf72* ALS/FTD iPSC-MG engulfing and degrading human synaptoneurosomes pHrodo (hSN-rodo)**

Here, we present selected time lapse live cell imaging of *C9orf72* ALS/FTD iPSC-MG engulfing hSN-rodo. Fluorescent live cell imaging with differential interference contrast microscopy was performed over a 6 h and 2 h time frame with images taken every 10 min. IPSC-MGs were identified using nuclear marker Hoechst (blue) followed by treatment with hSN-rodo (red). The increase in hSN-rodo intensity is observed in individual cells indicative of the uptake of hSN-rodo into acidic intracellular compartments for degradation.

